# *Dmp1Cre*-directed knockdown of PTHrP in murine decidua is associated with increased bone width and a life-long increase in strength specific to male progeny

**DOI:** 10.1101/2020.06.09.141812

**Authors:** Niloufar Ansari, Tsuyoshi Isojima, Blessing Crimeen-Irwin, Ingrid J Poulton, Narelle E. McGregor, Patricia W. M. Ho, Christopher S Kovacs, Evdokia Dimitriadis, Jonathan H Gooi, T. John Martin, Natalie A. Sims

## Abstract

Parathyroid hormone related-protein (PTHrP) is a pleiotropic regulator of tissue homeostasis. In bone, knockdown in osteocytes by *Dmp1Cre-*targeted deletion causes osteopenia and impaired strength. We report that this outcome depends on parental genotype. Adult *Dmp1Cre.Pthlh*^*f/f*^ mice from homozygous parents (*Dmp1Cre.Pthlh*^*f/f(hom)*^) have stronger bones, with 40% more trabecular bone mass and 30% greater femoral width than controls. At 12 days old, greater bone width was also found in male and female *Dmp1Cre.Pthlh*^*f/f(hom)*^ mice, but not in gene-matched mice from heterozygous parents, suggesting a maternal influence before weaning. Milk PTHrP levels were normal, but decidua from mothers of *Dmp1Cre.Pthlh*^*f/f(hom)*^ mice were smaller, with low PTHrP levels. Moreover, *Dmp1Cre.Pthlh*^*f/f(hom)*^ embryonic bone was more mineralized and wider than control. We conclude that *Dmp1Cre* leads to gene recombination in decidua, and that decidual PTHrP influences decidual cell maturation and limits embryonic bone growth. This identifies a maternal-derived developmental origin of adult bone strength.

## Introduction

Bone size and geometry are among the many factors determining bone strength (1). During skeletal development, bone grows in both the longitudinal and radial axes. Longitudinal growth is mediated by chondrocytes at the growth plates, where hypertrophic chondrocytes cease dividing, enlarge and eventually mineralize surrounding matrix (2). Simultaneously, expansion of bone diameter (termed “radial growth”) balances bone length and width. Although longitudinal growth has been studied widely, little is known about the signaling pathways orchestrating radial growth (3).

Parathyroid hormone-related protein (PTHrP, gene name: *Pthlh*) is produced by many tissues, and acts locally to maintain their physiological function (4). While global knockout of PTHrP is neonatal lethal and causes widespread skeletal defects including reduced bone length, due largely to PTHrP’s role in promoting chondrocyte maturation (5), heterozygous *Pthlh* deletion causes osteopenia in adult mice (6). Local PTHrP production by bone cells is also required for normal bone formation in adults during bone remodeling. This was established by studies in genetically altered mice; mice with *Pthlh* knockdown targeted to osteoblasts (*Col1(2.3kb)Cre.Pthlh*^*f/f*^) (6) or to osteocytes (*Dmp1(10kb)Cre*) (7) both exhibiting low bone formation and osteopenia in adulthood.

Here we report an effect of parental genotype on the bone structure of *Dmp1Cre.Pthlh*^*f/f*^ mice. This study arose from an unexpected finding when in follow up of our previous study (7), we sought to assess the effect of *Dmp1Cre-*targeted knockdown of PTHrP in osteocyte in older mice. For this, we changed our breeding strategy from using heterozygous breeders to homozygous breeders to limit mouse wastage. As in previous studies from our laboratory (8, 9), we generated these mice using cousin-bred homozygous breeding pairs. To our surprise, adult male PTHrP-deficient mice generated from homozygous breeders (denoted *Dmp1Cre.Pthlh*^*f/f(hom)*^) exhibited an opposing phenotype to that of mice used in our previous work, generated from heterozygous breeders (denoted *Dmp1Cre.Pthlh*^*f/f(het)*^) (7): adult male *Dmp1Cre.Pthlh*^*f/f(hom)*^ mice had high trabecular bone mass, and wide long bones, but normal body weight and normal bone length. Since this was a profound and reproducible phenotype, we sought to determine the parental source of the defect in bone structure.

Although we previously reported that *Dmp1Cre* can lead to gene recombination in the mammary gland (7), there was no alteration in milk PTHrP levels in *Dmp1Cre.Pthlh*^*f/f(het)*^ dams. However, suckling male and female *Dmp1Cre.Pthlh*^*f/f(hom)*^ mice both exhibited the wide bone phenotype. We traced the phenotype back to fetal development and found it was associated with low PTHrP levels in decidua basalis and impaired decidualization in mothers of *Dmp1Cre.Pthlh*^*f/f(hom)*^ mice. This implies that PTHrP from the decidua limits bone radial growth in male and female mice, and that this has life-long effects on skeletal size in males, that override the effects of endogenous PTHrP deletion in osteocytes. This has significant implications for bone development, sex-differences in bone growth, and for breeding strategies used with *Dmp1Cre-*targeted mouse models.

## Results

### Adult male *Dmp1Cre.Pthlh*^*f/f(hom)*^ mice have a high trabecular bone mass set-point, reached before 14 weeks of age

In contrast to our previous experiments showing osteopenia in 12 week old *Dmp1Cre.Pthlh*^*f/f(het)*^ mice (7), adult male *Dmp1Cre.Pthlh*^*f/f(hom)*^ mice (i.e. mice bred from parents expressing *Dmp1Cre* and homozygous for the *Pthlh*^*f/f*^ genotype, Figure 1A) had greater trabecular bone volume than age- and sex-matched controls (Figure 1B-G). At 14, 16 and 26 weeks of age, male *Dmp1Cre.Pthlh*^*f/f(hom)*^ mice had significantly higher trabecular bone volume (Figure 1B) and trabecular number (Figure 1C) than age- and sex-matched controls. This phenotype was stable; the proportional difference in trabecular bone volume and number between genotypes was the same at all three time points assessed: i.e. trabecular bone volume and number were at a constant ∼140% and ∼133% of sex-matched male controls. Trabecular thickness remained unchanged in *Dmp1Cre.Pthlh*^*f/f(hom)*^ mice compared to sex- and age-matched controls at all time points (Figure 1D). Trabecular separation was ∼18% lower than controls in male *Dmp1Cre.Pthlh*^*f/f(hom)*^ mice, and was statistically significant only at the age of 14 weeks (Figure 1E).

**Figure 1:**
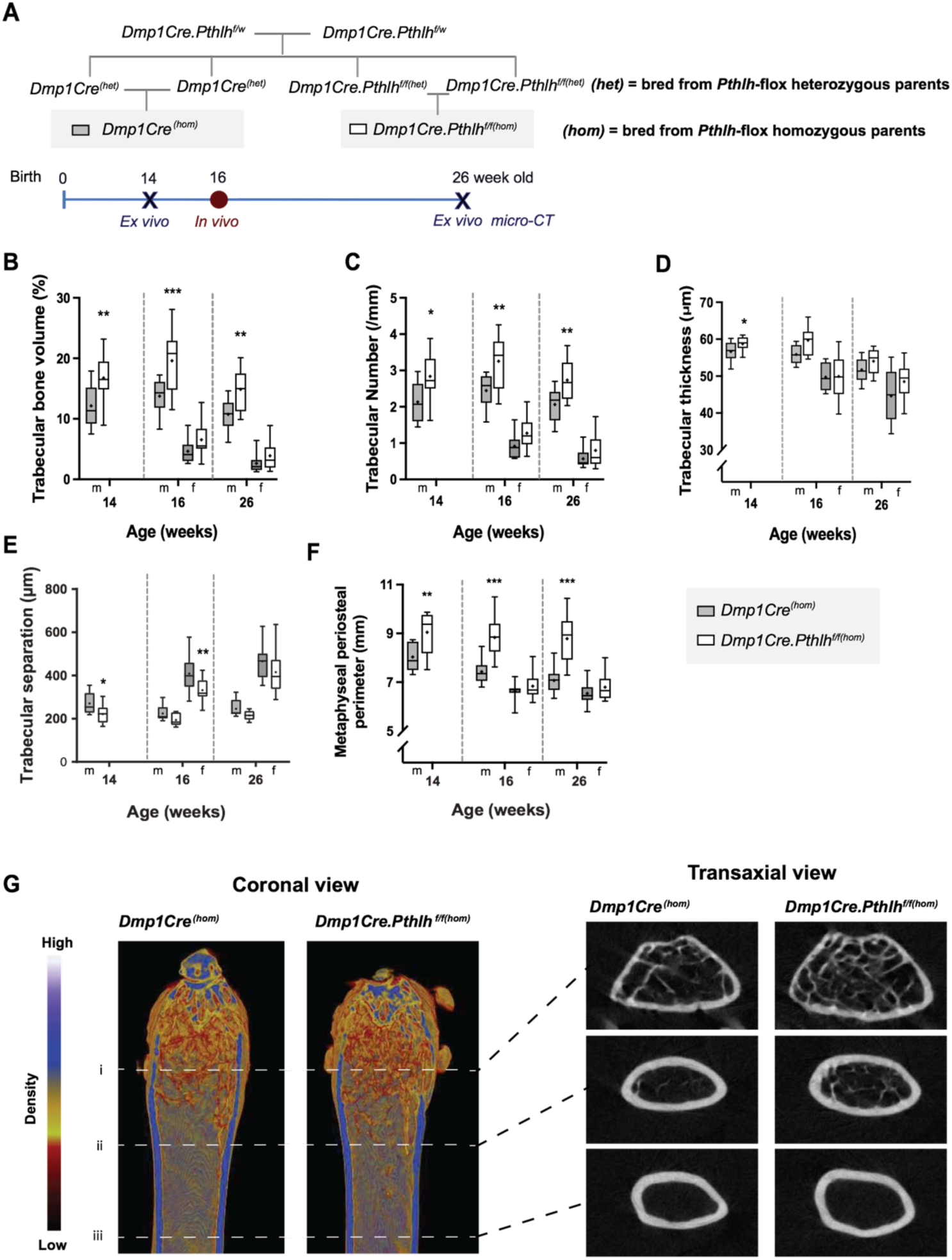
Breeding strategy and high trabecular bone mass in 14, 16 and 26 week old male *Dmp1Cre.Pthlh*^*f/f(hom)*^ mice. **A:** Schematic showing breeding strategy and data collection points. **B-F:** Trabecular structure of *Dmp1Cre.Pthlh*^*f/f(hom)*^ distal femoral primary spongiosa analysed by micro-CT in male mice (m) at 14, 16 and 26 weeks of age, and in female mice (f) at 16 and 26 weeks of age. Trabecular bone volume, trabecular number, trabecular thickness, trabecular separation, and metaphyseal periosteal perimeter are shown as mean (dot), interquartile range (box), median (line) and range; n=9-10/group. *p<0.05, **p<0.01, ***p<0.001 compared to sex- and age-matched *Dmp1Cre*^*(hom)*^ by two-way ANOVA (16 and 26 weeks old) and Student’s t-test (14 weeks old). **G**: Representative micro-CT images of trabecular bone in the distal femoral primary spongiosa of 26-week old male mice, showing density (scale above images), and raw cross-sectional images of the metaphysis (i), metaphyseal diaphysis (ii) and diaphysis (iii), showing a difference in bone size, and projection of trabecular bone into the lower metaphysis in *Dmp1Cre.Pthlh*^*f/f(hom)*^ samples.

When representative images were generated to show this difference in trabecular bone mass (Figure 1G), we noted that male *Dmp1Cre.Pthlh*^*f/f(hom)*^ mice also had wider bones; when measured in the trabecular region of analysis, metaphyseal periosteal perimeter was significantly higher in male *Dmp1Cre.Pthlh*^*f/f(hom)*^ mice at all three time points (Figure 1F), confirming the wider bone phenotype of these mice. When *Dmp1Cre.Pthlh*^*f/f(het)*^ mice were bred and aged to 16 and 26 weeks, no significant difference in trabecular structure or metaphyseal bone perimeter was detected at either age between *Dmp1Cre.Pthlh*^*f/f(het)*^ mice and their controls (Figure 1 Supplement 1). These indicate that male *Dmp1Cre.Pthlh*^*f/f(hom)*^ mice have a different phenotype to *Dmp1Cre.Pthlh*^*f/f(het)*^ mice (7), although they have the same genotype, showing that parental genotype influences trabecular bone mass and bone radial growth.

The high trabecular bone mass in *Dmp1Cre.Pthlh*^*f/f(hom)*^ mice was sex-specific. Female *Dmp1Cre.Pthlh*^*f/f(hom)*^ mice showed no significant difference in trabecular bone volume, trabecular number, or metaphyseal periosteal perimeter compared to *Dmp1Cre*^*(hom)*^ controls (Figure 1B-F). They did have lower trabecular separation at 16 weeks (Figure 1E), implying a mild and transient elevation in trabecular bone mass.

To confirm this trabecular phenotype in homozygous-bred mice, trabecular bone structure was studied at a second anatomical region, 5^th^ lumbar (L5) vertebrae. Similar to long bones, vertebrae of 14 week old male *Dmp1Cre.Pthlh*^*f/f(hom)*^ had higher trabecular bone volume and trabecular number than controls (Table 1). Trabecular separation was lower in male *Dmp1Cre.Pthlh*^*f/f(hom)*^ mice than controls, and there was no significant difference in trabecular thickness (Table 1). This confirmed the trabecular phenotype was not restricted to a single anatomical location.

**Table 1.** Trabecular structure of L5 (5^th^ lumbar vertebrae) from 14 week old male *Dmp1Cre.Pthlh*^*f/f(hom)*^ and *Dmp1Cre*^*(hom)*^ controls by microcomputed tomography. Values are mean ± SEM. n=9-10/group; * p<0.05 vs *Dmp1Cre*^*(hom)*^ by Student’s t-test.

|  | *Dmp1Cre<sup>(hom)</sup>* | *Dmp1Cre.Pthlh<sup>f/f(hom)</sup>* |
| --- | --- | --- |
| Trabecular bone volume/TV (%) | 17.84 ± 0.89 | **21.32 ± 1.16\*** |
| Trabecular number (/mm) | 3.20 ± 0.13 | **3.70 ± 0.15\*** |
| Trabecular thickness (μm) | 55.6 ± 1.3 | 57.3 ± 1.0 |
| Trabecular separation (μm) | 203.5 ± 4.9 | **185.4 ± 6.4\*** |

Although trabecular bone mass was greater in male 14 week old *Dmp1Cre.Pthlh*^*f/f(hom)*^ mice, dynamic histomorphometry revealed no difference in any bone formation or resorption parameters compared to controls (Table 2). In addition, no significant difference was detected in serum levels of the bone formation or resorption markers, P1NP and CTX1, of 14 week old male *Dmp1Cre.Pthlh*^*f/f(hom)*^ mice compared to controls (Table 2). This, and the similar proportion of elevation in trabecular bone mass at 14, 16 and 26 weeks suggest that the high trabecular bone mass arose before 14 weeks of age, and has reached a greater adult set-point for “peak bone mass” than controls.

**Table 2.** Histomorphometry of tibia and serum biochemical data of 14 week-old male *Dmp1Cre.Pthlh*^*f/f(hom)*^ and *Dmp1Cre*^*(hom)*^ controls. Histomorphometry was measured in the distal tibial metaphyseal secondary spongiosa. Values are mean ± SEM. n=7-10/group. Abbreviations: BV: bone volume; BS: bone surface; B. Pm: bone perimeter; N.Ob: osteoblast numbers; B.Pm: bone perimeter; N.Oc: osteoclast numbers; Oc.Pm: osteoclast perimeter; P1NP: Procollagen type 1 N-terminal propeptide; CTX1: C-telopeptide of type 1 collagen.

|  | *Dmp1Cre<sup>(hom)</sup>* | *Dmp1Cre.Pthlh<sup>f/f(hom)</sup>* |
| --- | --- | --- |
| Osteoid surface/BS (%) | 6.16 ± 1.85 | 6.47 ± 1.83 |
| Osteoid thickness (μm) | 0.90 ± 0.11 | 1.20 ± 0.23 |
| Osteoblast surface/BS (%) | 7.83 ± 2.46 | 9.02 ± 2.55 |
| Osteoblast numbers (N.Ob/B.Pm) (N/mm) | 6.72 ± 2.09 | 7.77 ± 2.16 |
| Osteoclast surface/BS (%) | 3.77 ± 0.38 | 3.86 ± 0.44 |
| Osteoclast numbers (N.Oc/B.Pm) (N/mm) | 1.79 ± 0.29 | 1.93 ± 0.23 |
| N.Oc/Oc.Pm (N/mm) | 46.60 ± 6.18 | 49.79 ± 1.50 |
| Single-labeled mineralizing surface (sL.S/BS) (%) | 22.4 ± 2.4 | 20.8 ± 3.8 |
| Double-labeled mineralizing surface (dL.S/BS) (%) | 12.9 ± 2.9 | 15.9 ± 4.2 |
| Mineralising surface/BS (%) | 24.2 ± 2.0 | 26.3 ± 3.3 |
| Mineral apposition rate (μm/day) | 1.09 ± 0.08 | 1.05 ± 0.15 |
| Bone formation rate/BV (%/day) | 1.30 ± 0.10 | 1.41 ± 0.32 |
| P1NP (ng/ml) | 29.6 ± 2.3 | 36.6 ± 4.1 |
| CTX1 (ng/ml) | 25.4 ± 1.8 | 21.5 ± 1.6 |

### Adult male *Dmp1Cre.Pthlh*^*f/f(hom)*^ mice have a wide bone phenotype, reached before 14 weeks of age, that leads to greater bone strength

Since the metaphyseal bone width was greater in adult male *Dmp1Cre.Pthlh*^*f/f(hom)*^ mice than controls, we analysed femoral cortical bone structure in more detail (Figure 2A). Although femoral length was not different between *Dmp1Cre.Pthlh*^*f/f(hom)*^ mice and *Dmp1Cre*^*(hom)*^ controls (Figure 2B), adult male *Dmp1Cre.Pthlh*^*f/f(hom)*^ femora were wider in both anteroposterior and mediolateral dimensions compared to controls at 14, 16 and 26 weeks (Figure 2C,D). The greater femoral width in male *Dmp1Cre.Pthlh*^*f/f(hom)*^ mice was in proportion: the ratio of anteroposterior to mediolateral widths was not different in these mice compared to controls at any time point (data not shown). Adult male *Dmp1Cre.Pthlh*^*f/f(hom)*^ mice exhibited greater marrow and cortical area, and greater periosteal and endocortical perimeters, compared to age- and sex-matched controls (Figure 2E-I). Although cortical diameter was greater, this was balanced on the endocortical and periosteal surfaces, as there was no significant difference in cortical thickness (Figure 2G). As in trabecular bone, the greater cortical bone width phenotype was stable, showing a similar proportional difference compared to controls at all three time points.

**Figure 2:**
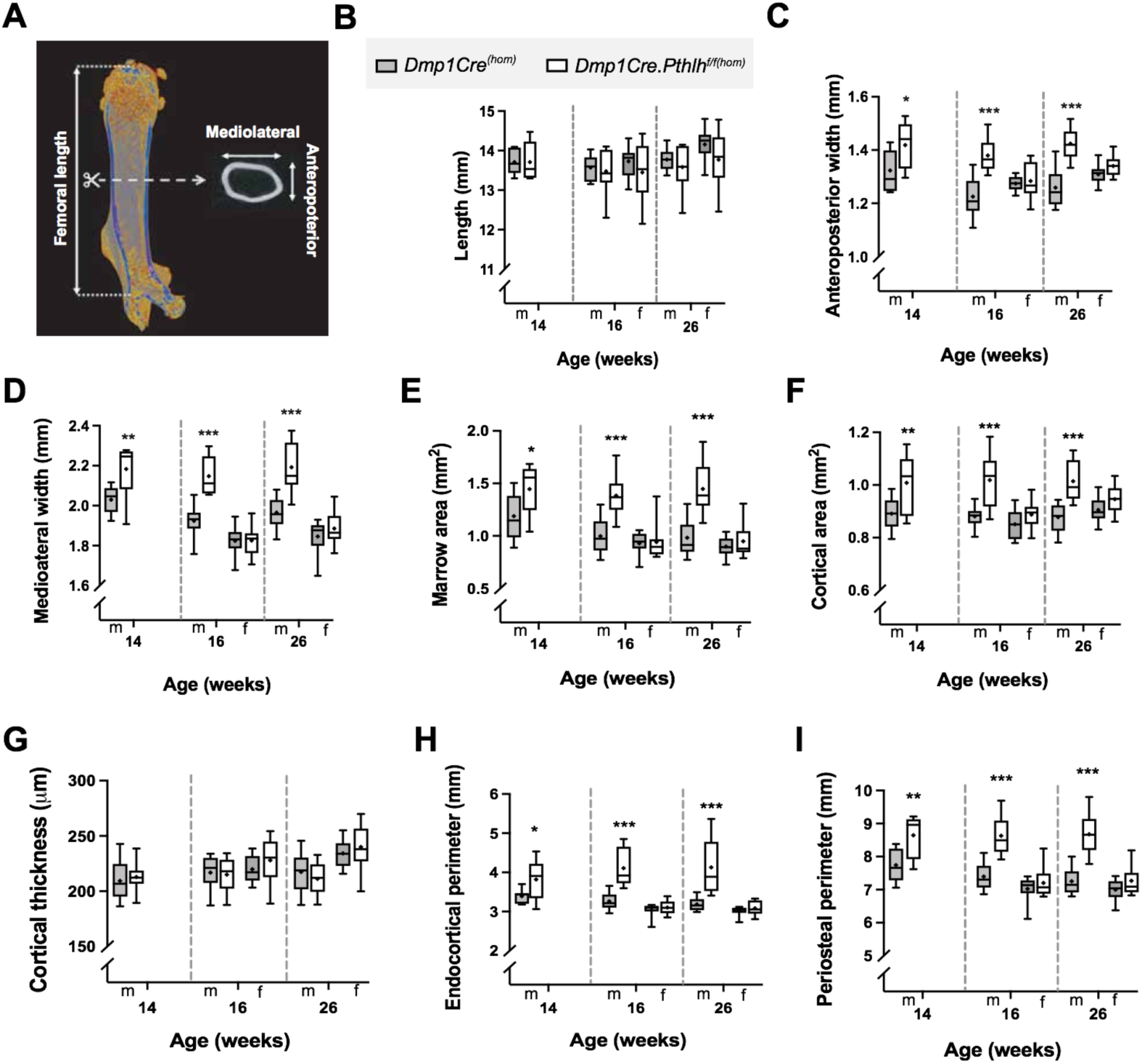
Wide bone phenotype of adult male *Dmp1Cre.Pthlh*^*f/f(hom)*^ mice. **A**: Schematic showing measurement regions. Bone length (**B**) and cortical dimensions of male (m) *Dmp1Cre.Pthlh*^*f/f(hom)*^ and *Dmp1Cre*^*(hom)*^ mice at 14, 16 and 26 weeks of age, and female mice (f) at 16 and 26 weeks of age. Anteroposterior (**C**) and mediolateral (**D**) width, measured by micro-CT at the midshaft. **E:I** Femoral marrow area (**E**), cortical bone area (**F**), thickness (**G**), and both endocortical (**H**) and periosteal (**I**) perimeter were analysed in cortical ROI by micro-CT. Data are shown as mean (dot), interquartile range (box), median (line) and range; n=9-10/group. *p<0.05, **p<0.01, ***p<0.001 compared to sex- and age-matched *Dmp1Cre*^*(hom)*^ by two-way ANOVA (16 and 26 weeks old) and Student’s t-test (14 weeks old).

Female *Dmp1Cre.Pthlh*^*f/f(hom)*^ mice showed no significant difference in cortical bone size or shape compared to age-matched *Dmp1Cre*^*(hom)*^ mice at 16 or 26 weeks of age (Figure 2).

Heterozygous-bred *Dmp1Cre.Pthlh*^*f/f(het)*^ mice at 16 and 26 weeks of age did not exhibit any significant difference in anteroposterior or mediolateral femoral width compared to *Dmp1Cre*^*(het)*^ controls (Figure 2 Supplement 1B,C), nor in marrow area, or periosteal circumference (Figure 2 Supplement 1E,I). At 16 weeks, male mice did exhibit a small and significant transient elevation in cortical area and endocortical perimeter; this was no longer detected at 26 weeks (Figure 2 Supplement 1F,H).

Since greater bone width is associated with greater bone strength, we carried out three-point bending tests. Femora from 26 week old male *Dmp1Cre.Pthlh*^*f/f(hom)*^ mice could withstand higher loads than age-matched *Dmp1Cre*^*(hom)*^ controls, reaching a higher ultimate force (Figure 3A,B) and failure force (Table 3) before breaking. There was no significant difference in ultimate displacement between *Dmp1Cre.Pthlh*^*f/f(hom)*^ and control femora (Figure 3C). When these measurements were corrected for bone size, both 14 and 26 week old male *Dmp1Cre.Pthlh*^*f/f(hom)*^ femora showed lower ultimate stress and yield stress, compared to controls (Figure 3D-F). 14 week old male *Dmp1Cre.Pthlh*^*f/f(hom)*^ femora had higher ultimate and failure strain than controls (Table 3). 26 week old male *Dmp1Cre.Pthlh*^*f/f(hom)*^ femora had lower failure stress (Figure 3G), and reduced toughness and elastic modulus (Table 3) compared to sex- and age-matched controls. This indicates that the greater width of the bones increased bone strength by lowering the stress experienced by the material under three point bending conditions.

**Table 3.** Additional strength data from three point bending test on femora from 14 and 26 week old *Dmp1Cre.Pthlh*^*f/f(hom)*^ (f/f) and *Dmp1Cre*^*(hom)*^ (w/w) mice. Values are mean ± SEM. n=9-10/group; *p<0.05, **p<0.01, and ***p<0.001 vs sex- and age-matched w/w by Student’s t-test (14 weeks old) and two-way ANOVA (26 weeks old) with uncorrected Fishers LSD post-hoc test.

|  | 14 week old |  | 26 week old |  |  |  |
| --- | --- | --- | --- | --- | --- | --- |
|  | Males |  | Males |  | Females |  |
|  | w/w | f/f | w/w | f/f | w/w | f/f |
| Moment of inertia (mm<sup>4</sup>) | 0.76 ± 0.06 | **1.16 ± 0.10\*\*** | 0.62 ± 0.05 | **1.16 ± 0.09\*\*\*** | 0.59 ± 0.03 | 0.67 ± 0.03 |
| **Whole bone parameters** |  |  |  |  |  |  |
| Yield force (N) | 12.64 ± 0.63 | 13.85 ± 0.94 | 11.50 ± 0.44 | 13.71 ± 0.96 | 12.61 ± 0.84 | **15.73 ± 1.61\*** |
| Yield displacement (μm) | 436.9 ± 56.2 | 454.4 ± 43.2 | 157.4 ± 13.5 | 161.3 ± 15.2 | 132.4 ± 11.5 | **190.8 ± 25.5\*** |
| Post-yield displacement (μm) | 298.4 ± 53.6 | 388.9 ± 44.4 | 293.7 ± 20.1 | 238.3 ± 36.9 | 200.0 ± 38.9 | 154.2 ± 28.9 |
| Failure force (N) | 14.26 ± 1.63 | 15.88 ± 1.21 | 14.99 ± 0.47 | **18.25 ± 0.73\*** | 18.04 ± 0.97 | 20.06 ± 1.18 |
| Stiffness (N/mm) | 49.8 ± 7.0 | 66.9 ± 7.3 | 92.33 ± 4.95 | 111.06 ± 9.77 | 114.02 ± 4.07 | 106.27 ± 6.93 |
| **Tissue level parameters** |  |  |  |  |  |  |
| Ultimate strain (%) | 5.31 ± 0.46 | **6.81 ± 0.47\*** | 6.84 ± 0.45 | 8.13 ± 0.85 | 5.97 ± 0.29 | 7.17 ± 0.50 |
| Yield strain (%) | 4.23 ± 0.50 | 4.77 ± 0.46 | 3.30 ± 0.28 | 3.81 ± 0.34 | 2.88 ± 0.24 | **4.25 ± 0.55\*** |
| Post-yield strain (%) | 2.91 ± 0.50 | 4.07 ± 0.46 | 6.15 ± 0.41 | 5.69 ± 0.92 | 4.38 ± 0.86 | 3.43 ± 0.64 |
| Failure strain (%) | 7.15 ± 0.29 | **8.84 ± 0.54\*\*** | 9.44 ± 0.40 | 9.50 ± 1.02 | 7.26 ± 0.75 | 7.68 ± 0.52 |
| Toughness (mJ/mm<sup>3</sup>) | 1.33 ± 0.10 | 1.29 ± 0.11 | 1.74 ± 0.13 | **1.13 ± 0.13\*\*** | 1.53 ± 0.19 | 1.42 ± 0.13 |
| Elastic modulus (MPa) | 1009.4 ± 136.4 | 852.4 ± 73.2 | 692.1 ± 48.4 | **450.0 ± 47.2\*\*** | 891.5 ± 57.2 | **718.9 ± 49.5\*** |

**Figure 3:**
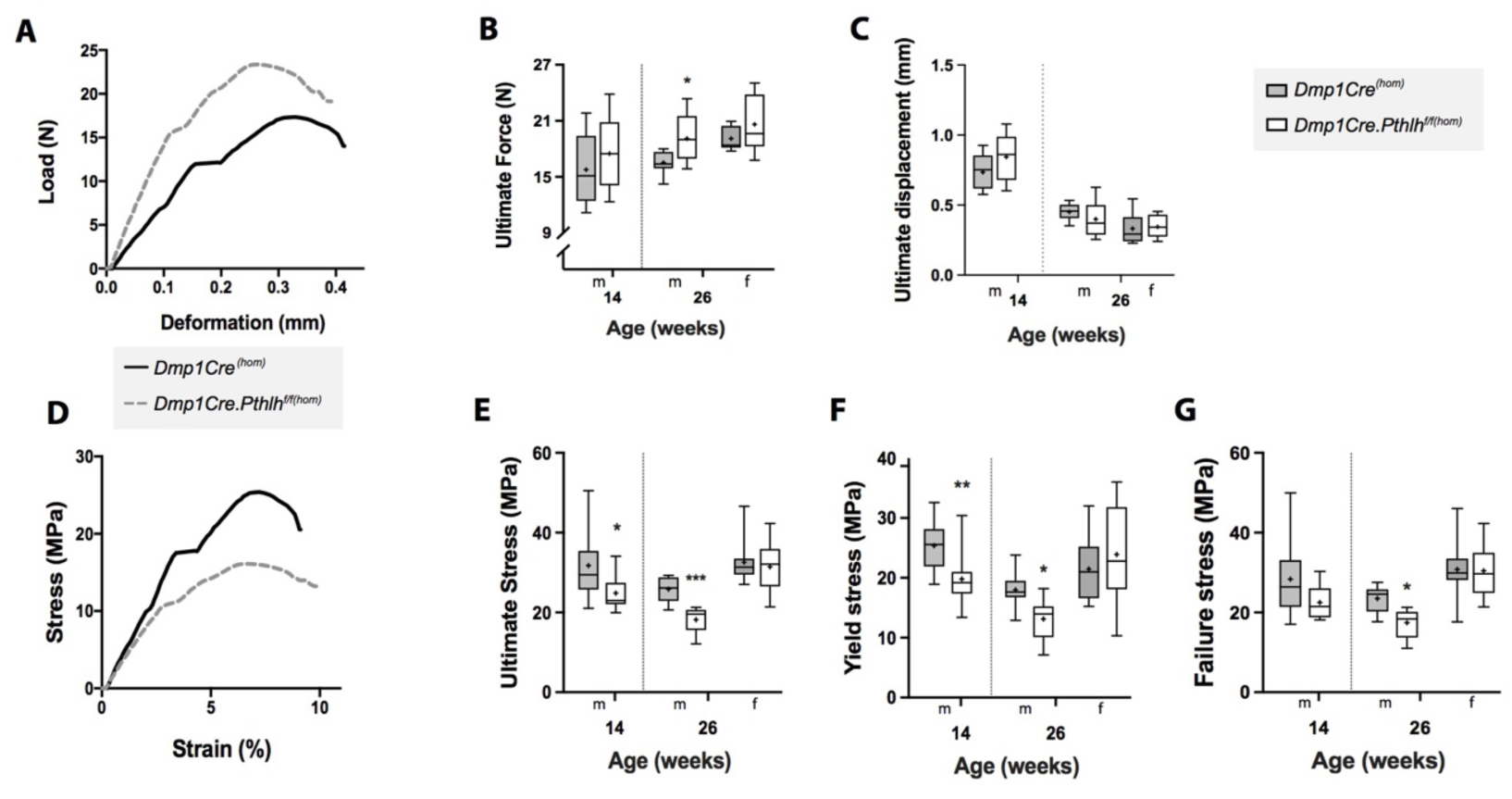
Greater ultimate strength in 26-week old and less stress in 14 and 26-week old male *Dmp1Cre.Pthlh*^*f/f(hom)*^ femora measured by three-point bending tests. **A:** Representative load-deformation curves of 26-week old male *Dmp1Cre.Pthlh*^*f/f(hom)*^ and *Dmp1Cre*^*(hom)*^ bones. Greater ultimate force (**B**) and normal ultimate deformation (**C**) in femora from 26-week old males; no change in 14 week old males and 26 week old females. **D**: Representative stress-strain curves of 26-week old male *Dmp1Cre.Pthlh*^*f/f(hom)*^ and *Dmp1Cre*^*(hom)*^ bones, after correction in each sample based on anteroposterior and mediolateral dimensions (shown in Figure 2). Also shown are ultimate stress (**E**), yield stress (**F**), and failure stress (**G**). Data shown as mean (dot), interquartile range (box), median (line) and range, n=9-10/group. *p<0.05, **p<0.01 and ***p<0.001 compared to age- and sex-matched controls by two-way ANOVA (16 and 26 weeks old) and Student’s t-test (14 weeks old).

While ultimate force, failure force, ultimate stress, yield stress, failure stress and toughness were modified in the male mice, these parameters were not changed in females (Figure 3 and Table 3), consistent with their loss of the greater bone width with ageing. Surisingly, femora from female *Dmp1Cre.Pthlh*^*f/f(hom)*^ mice achieved a greater yield force and displacement compared to age- and sex-matched controls (Table 3), suggesting a higher elastic deformation. When corrected for bone size, 26 week old female *Dmp1Cre.Pthlh*^*f/f(hom)*^ femora had higher yield strain and lower elastic modulus (Table 3), suggesting a more flexible material than controls.

Femora from 26 week old heterozygous-bred *Dmp1Cre.Pthlh*^*f/f(het)*^ mice did not show any significant difference in mechanical properties compared to *Dmp1Cre*^*(het)*^ controls (Figure 3 Supplement 1).

### The wide bone phenotype is present at 12 days of age in male and female *Dmp1Cre.Pthlh*^*f/f(hom)*^ mice

Since we previously observed recombination of PTHrP in the mammary gland in *Dmp1Cre.Pthlh*^*f/f(het)*^ mice (7), and deletion of maternal mammary PTHrP is reported to lead to greater bone mass in progeny at 12 days of age (10), we sought to determine whether bone mass was modified in 12 day old *Dmp1Cre.Pthlh*^*f/f(hom)*^ mice, and whether maternal milk PTHrP content was reduced. 12 day old mice were assessed from both heterozygous (*Dmp1Cre.Pthlh*^*f/f(het)*^) (Figure 4A) and homozygous (*Dmp1Cre.Pthlh*^*f/f(hom)*^) (Figure 1A) breeding strategies.

**Figure 4:**
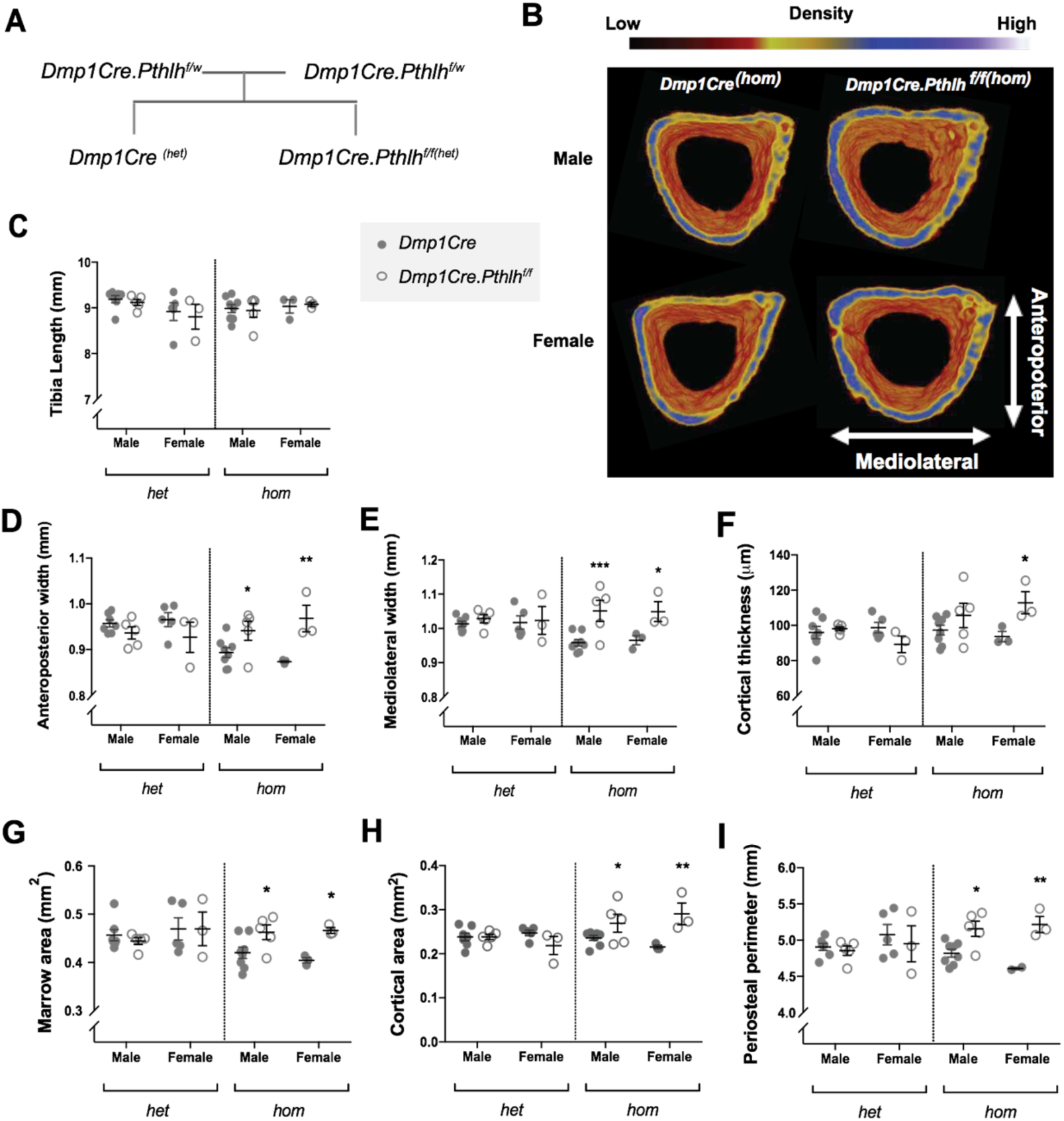
Wide bone phenotype of 12 day old male and female *Dmp1Cre.Pthlh*^*f/f(hom)*^ mice. **A:** Schematic diagram showing heterozygous breeding strategy used for this experiment. **B:** Representative micro-CT images of cortical bone of 12 day-old *Dmp1Cre.Pthlh*^*f/f(hom)*^ tibiae. **C-I:** Tibial cortical structure of 12 day old *Dmp1Cre.Pthlh*^*f/f*^ mice from heterozygous (het) and homozygous (hom) breeders compared to their respective *Dmp1Cre* controls. Shown are tibial length (**C**), anteroposterior (**D**) and mediolateral (**E**) width, cortical thickness (**F**), marrow area (**G**), cortical area (**H**) and periosteal perimeter (**I**). Data is shown as mean ± SEM with individual data points, *p<0.01, **p<0.01, ***p<0.001 compared to age- and sex-matched controls by two-way ANOVA.

While no significant differences in cortical dimensions were detected in tibiae from male or female *Dmp1Cre.Pthlh*^*f/f(het)*^ mice (from heterozygous breeders), *Dmp1Cre.Pthlh*^*f/f(hom)*^ mice (from homozygous breeders) exhibited greater tibial width at 12 days of age (Figure 4B-I) with no difference in tibial length (Figure 4C). Both male and female *Dmp1Cre.Pthlh*^*f/f(hom)*^ mice had significantly greater tibial width in the anteroposterior and mediolateral direction, compared to sex-matched *Dmp1Cre*^*(hom)*^ (Figure 4B,D,E). Male and female *Dmp1Cre.Pthlh*^*f/f(hom)*^ mice also showed greater tibial marrow area (Figure 4G), cortical area (Figure 4H), and periosteal perimeter (Figure 4I) compared to sex-matched cousin-bred *Dmp1Cre*^*(hom)*^ controls. Female *Dmp1Cre.Pthlh*^*f/f(hom)*^ also had greater cortical thickness than female *Dmp1Cre*^*(hom)*^ controls (Figure 2F). No significant difference was observed in trabecular structure between *Dmp1Cre.Pthlh*^*f/f(hom)*^ mice and their *Dmp1Cre*^*(hom)*^ sex-matched controls (Table 4), suggesting that this aspect of the phenotype was secondary to the increase in bone width.

**Table 4.** Trabecular bone structure of 12 day old *Dmp1Cre.Pthlh*^*f/f(hom)*^ and *Dmp1Cre*^*(hom)*^ mice. Trabecular bone was measured by histomorphometry in male and female tibiae. Data is shown as mean ± SEM.

|  | Male |  | Female |  |
| --- | --- | --- | --- | --- |
|  | *Dmp1Cre<sup>(hom)</sup>* (n=6) | *Dmp1Cre.Pthlh<sup>ff(hom)</sup>*<br>(n=4) | *Dmp1Cre<sup>(hom)</sup>*<br>(n=4) | *Dmp1Cre.Pthlh<sup>ff(hom)</sup>* (n=3) |
| Trabecular bone volume (%) | 3.65 ± 1.00 | 5.19 ± 2.11 | 2.78 ± 0.98 | 8.38 ± 4.70 |
| Trabecular number (/mm) | 1.36 ± 0.30 | 1.84 ± 0.44 | 1.12 ± 0.32 | 2.51 ± 0.92 |
| Trabecular thickness (μm) | 24.85 ± 2.38 | 25.63 ± 3.78 | 24.13 ± 2.27 | 29.15 ± 5.97 |
| Trabecular separation (μm) | 1101.06 ± 407.40 | 589.18 ± 103.76 | 1344.61 ± 607.52 | 476.28 ± 155.84 |

Although PTHrP gene recombination was detected in mammary tissue from *Dmp1Cre.Pthlh*^*f/f(het)*^ mice (7), milk PTHrP levels, measured either by radioimmunoassay or bioassay, and milk protein levels were not significantly altered in *Dmp1Cre.Pthlh*^*f/f(het)*^ mice compared to controls (Table 5).

**Table 5.**
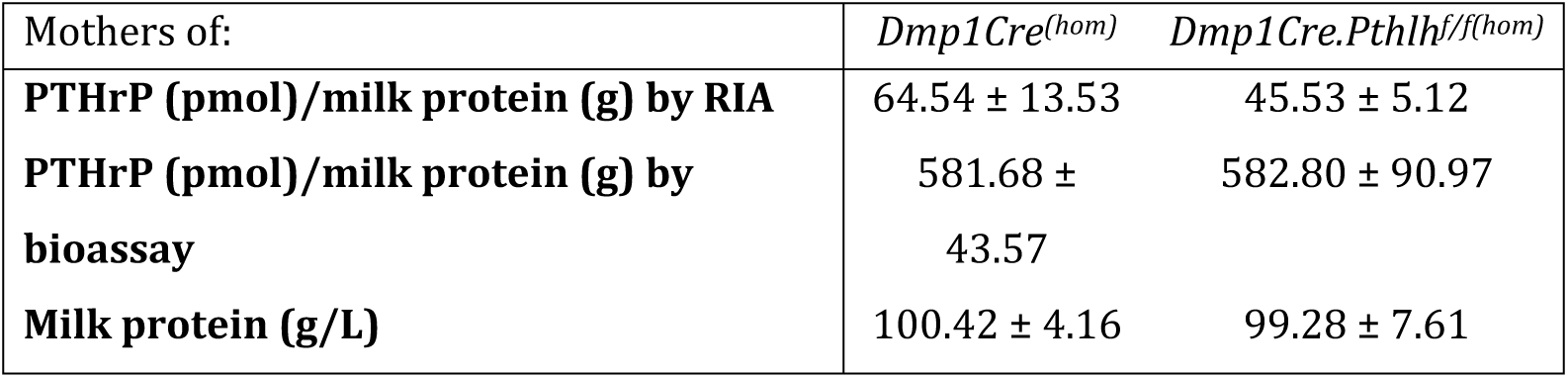
Biochemical analysis of milk samples. PTHrP content was measured by radioimmunoassay (RIA) in milk samples from mothers of *Dmp1Cre.Pthlh*^*f/f(hom)*^ and *Dmp1Cre*^*(hom)*^ mice on day 12 of lactation. PTHrP equivalent concentration was measured by measuring cAMP response to milk treatment of UMR106-01 cells (bioassay). Data shown as mean ± SEM. n=7/group. No significant differences relating to genotype were detected by Student’s t-test.

### The *Dmp1Cre.Pthlh*^*f/f(hom)*^ wide-bone phenotype exists *in utero*

Since no change in milk PTHrP could explain the phenotype at 12 days of age, we determined whether the phenotype existed *in utero* by assessing embryonic bone size. Consistent with our observations at 12 days, embryonic *Dmp1Cre.Pthlh*^*f/f(hom)*^ femora (E18.5) were wider in both anteroposterior and mediolateral dimensions, and exhibited a higher moment of inertia compared to *Dmp1Cre*^*(hom)*^ controls (Figure 5A, D-I). Greater bone area and tissue area of *Dmp1Cre.Pthlh*^*f/f(hom)*^ embryos confirmed that their femora are wider than controls (Figure 5 Supplement 1). This greater bone width was observed along the full length of the bone, including both metaphysis and the diaphysis.

**Figure 5:**
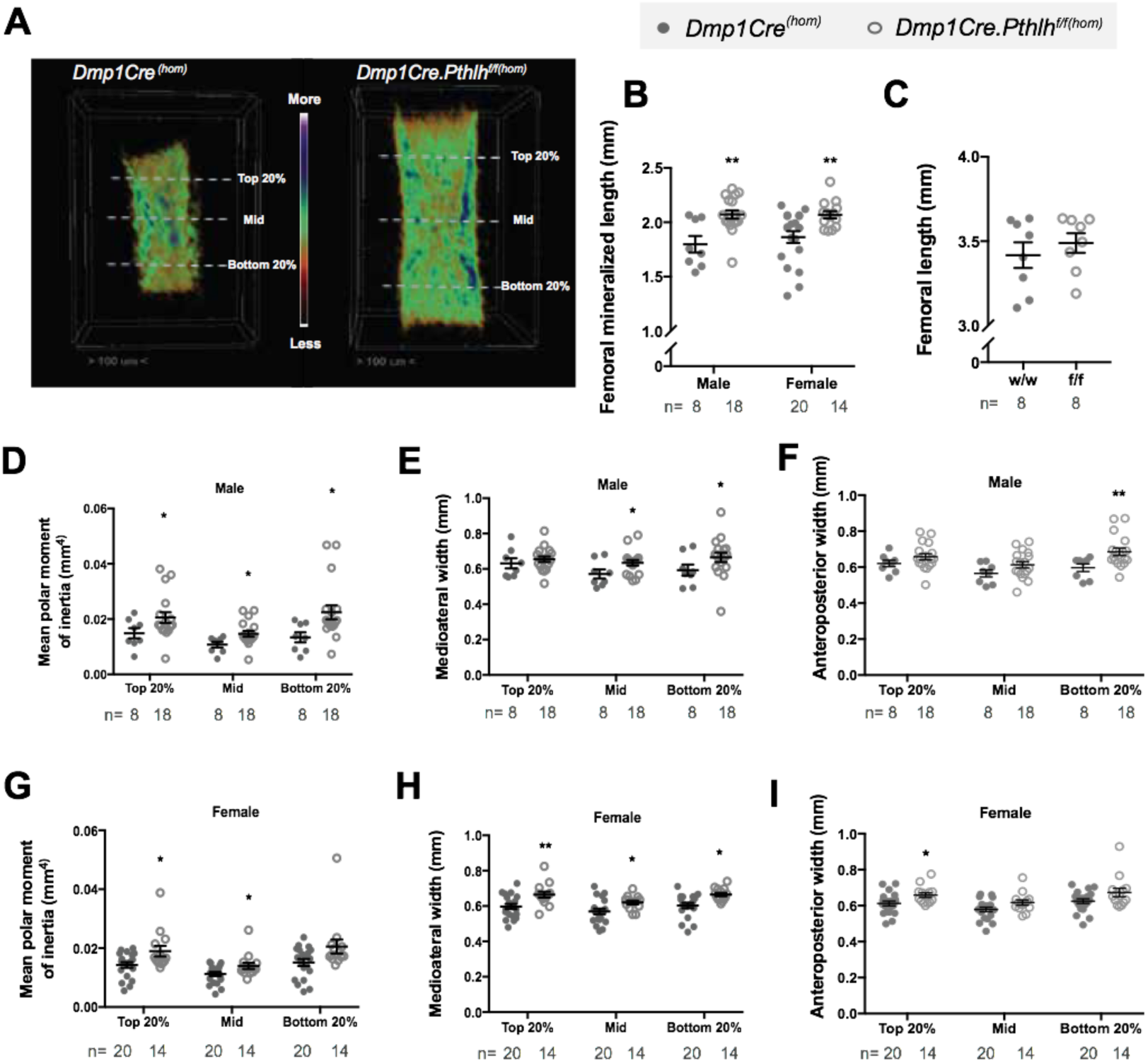
Accelerated bone development of E18.5 *Dmp1Cre.Pthlh*^*f/f(hom)*^ femora. **A**) representative images of *Dmp1Cre.Pthlh*^*f/f(hom)*^ and *Dmp1Cre*^*(hom)*^ femora at E18.5, showing bone mineral density. **B**) Femoral mineralized length, **C)** total femoral length, **D-I:** Mean polar moment of inertia (**D, G**), mediolateral (**E, H**), and anteroposterior widths (**F, I**) of *Dmp1Cre.Pthlh*^*f/f(hom)*^ and *Dmp1Cre*^*(hom)*^ at three different locations shown in panel A: at 20% of the mineralized length distal to the proximal end of the mineralized region (Top 20%), at the midshaft (Mid), and at 20% of the mineralized length proximal to the distal end of the mineralized region (Bottom 20%). Data is shown as mean ± SEM with individual data points, *p<0.05, **p<0.01, and ***p<0.001 compared to controls by two-way ANOVA (B-H).

The length of the mineralized portion of *Dmp1Cre.Pthlh*^*f/f(hom)*^ femora, measured by micro-CT, was also greater than *Dmp1Cre*^*(hom)*^ controls (Figure 5B). Micro-computed tomography cannot detect the full length of the bone at this age, as the cartilage ends are not yet mineralized prior to birth. Total femoral length measured in Alizarin Red/Alcian Blue stained samples showed no difference between *Dmp1Cre.Pthlh*^*f/f(hom)*^ and *Dmp1Cre*^*(hom)*^ controls (Figure 5C), indicating that *Dmp1Cre.Pthlhf/f*^*(hom)*^ embryos have normal bone length, but accelerated mineralization *in utero*.

Heterozygous-bred *Dmp1Cre.Pthlh*^*f/f(het)*^ embryos did not show any significant difference in the mineralized length compared to *Dmp1Cre*^*(het)*^ controls (Figure 5 Supplement 2A,B), nor in the bone width or structure (Figure 5 Supplement 2C-K).

### Low PTHrP levels and modified cell morphology in *Dmp1Cre.Pthlh*^*f/f*^ decidua

Since we observed increased bone width *in utero* in *Dmp1Cre.Pthlh*^*f/f(hom)*^ mice, and PTHrP is produced by uterus and decidua (11, 12), we sought to determine whether PTHrP expression is modified in placenta or decidua from *Dmp1Cre.Pthlh*^*f/f(hom)*^ mice. Consistent with a lack of change in overall bone growth, there were no significant differences in body weight, placental weight, or body to placental weight ratios between *Dmp1Cre.Pthlh*^*f/f(hom)*^ embryos and *Dmp1Cre*^*(hom)*^ controls (Figure 6A-C). No alteration in these parameters were observed in *Dmp1Cre.Pthlh*^*f/f(het)*^ embryos and placenta compared to littermate *Dmp1Cre.Pthlh*^*w/w(het)*^ (Figure 6 Supplement 1A-D). This suggests the increased bone width of *Dmp1Cre.Pthlh*^*f/f(hom)*^ embryos is not caused by changes in placental efficiency.

**Figure 6:**
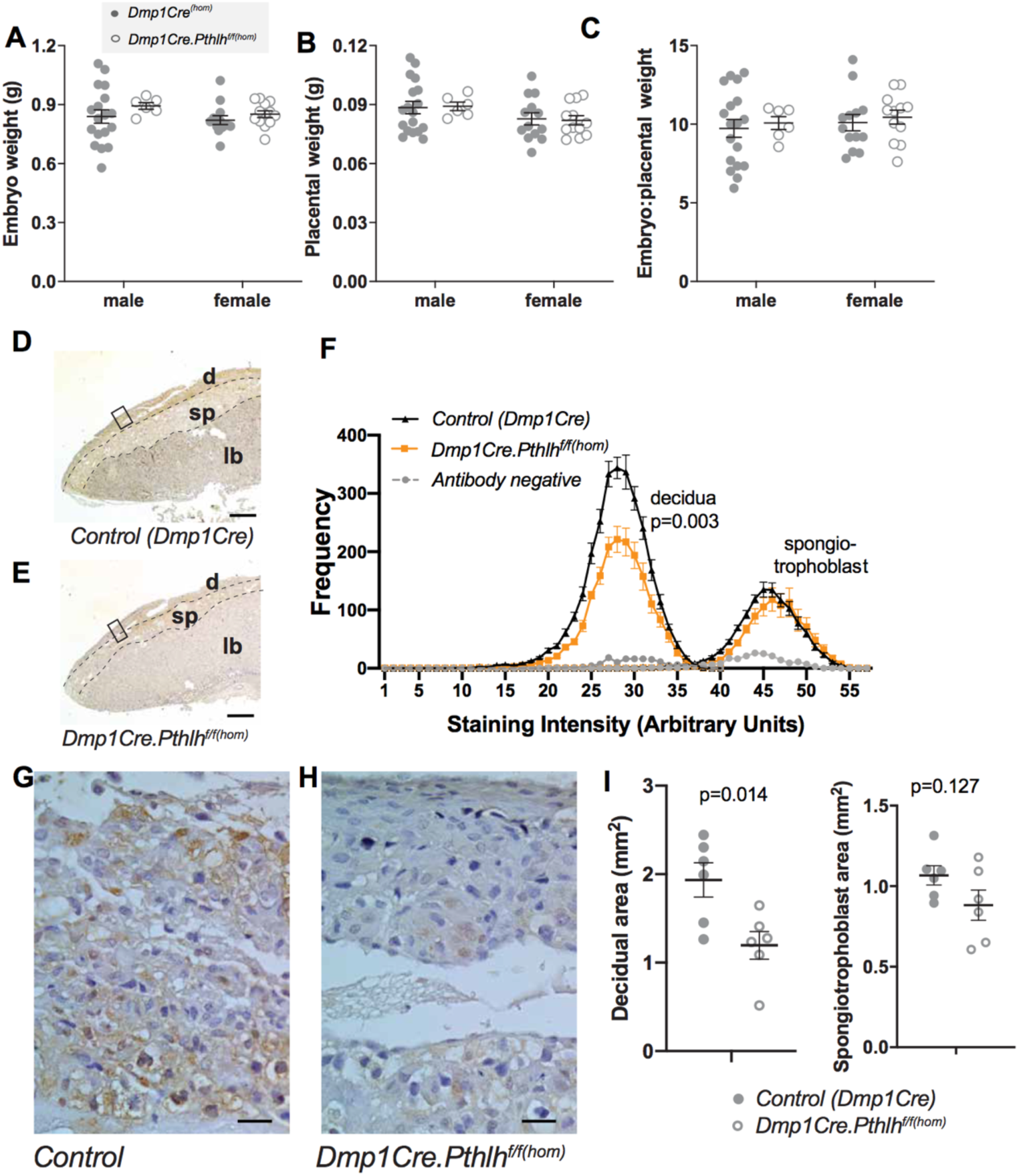
Decreased decidual PTHrP and impaired decidualization of *Dmp1Cre.Pthlh*^*f/f*^ mice at E17.5. Embryo weight **(A)**, placental weight **(B)** and embryo to placental weight ratio **(C)** of *Dmp1Cre.Pthlh*^*f/f(hom)*^ and *Dmp1Cre*^*(hom)*^. **(D-H)** Immunostaining for PTHrP in samples of placenta and decidua from *Dmp1Cre.Pthlh*^*f/f(hom)*^ and *Dmp1Cre*^*(hom)*^ embryos at E17.5; IgG control staining was measured in images of both zones. Decidua (d), spongiotrophoblast (sp) and labyrinth (lb) zones are shown in low power images (D,E). Scale bar = 1 mm. Frequency of PTHrP stained objects segregated by staining intensity in the spongiotrophoblast layer and decidua from *Dmp1Cre.Pthlh*^*f/f(hom)*^ and *Dmp1Cre*^*(hom)*^ embryos. **(F**,**G)** High power images of decidua. Scale bar = 20 micron. **(I)** Quantitation of total decidual area and spongiotrophoblast area; mean ± SEM with individual data points, *p<0.05, **p<0.01compared to controls by one-way ANOVA.

PTHrP staining of decidua and placenta showed positive staining for PTHrP in both the decidua and the spongiotrophoblast layer (junctional zone) of the placenta (Figure 6D). No PTHrP was detected in the placental labyrinth zone. PTHrP staining in decidua from mothers of *Dmp1Cre.Pthlh*^*f/f(hom)*^ mice was not as strong as that observed in decidua from mothers of *Dmp1Cre*^*(hom)*^ mice (Figure 6D,E). Quantification revealed a significant reduction in PTHrP staining at all intensities in decidua from mothers of *Dmp1Cre.Pthlh*^*f/f(hom)*^ mice, but no change in PTHrP staining frequency in the spongiotrophoblast zone of the adjacent placenta (Figure 6F). IgG isotype control had minimal intensity in both regions. No alteration in PTHrP staining frequency was observed in in decidua adjacent to *Dmp1Cre.Pthlh*^*f/f(het)*^ placenta compared to littermate *Dmp1Cre.Pthlh*^*w/w(het)*^ (Figure 6 Supplement 1E). This suggests off-target effects of *Dmp1Cre* have led to reduced PTHrP protein production by decidual cells.

We also examined the morphology of decidua from samples adjacent to *Dmp1Cre.Pthlh*^*f/f(hom)*^ placenta. The decidual cells from mothers of *Dmp1Cre.Pthlh*^*f/f(hom)*^ embryos appeared more compact than in *Dmp1Cre*^*(hom)*^ decidua, suggesting impaired decidual cell maturation (Figure 6G,H). Total decidual area was significantly less in samples from mothers of *Dmp1Cre.Pthlh*^*f/f(hom)*^ embryos than *Dmp1Cre*^*(hom)*^, but the area of the spongiotrophoblast zone was not significantly modified (Figure 6I). No change in decidual size was detected in decidua adjacent to *Dmp1Cre.Pthlh*^*f/f(het)*^ placenta compared to littermate *Dmp1Cre.Pthlh*^*w/w(het)*^ controls (Figure 6 Supplement 1F,G). This suggests PTHrP may act locally within the decidua to promote decidual cell maturation, and this may cause increased bone width growth of *Dmp1Cre.Pthlh*^*f/f(hom)*^ mice *in utero*.

## Discussion

This study identifies that off-target effects of *Dmp1Cre*-mediated recombination led to reduced decidual PTHrP. Reduced PTHrP level in the decidua is associated with increased embryonic bone radial growth and mineralization *in utero*. This wide bone phenotype is observed in both male and females at 12 days of age, and is sustained until at least 6 months of age in male, but not female, skeletons (Figure 7). These effects of reduced decidual PTHrP on bone size, trabecular bone mass, and bone strength dominates over the previously reported effects of reducing endogenous PTHrP in osteocytes, which suppressed bone formation and reduced trabecular bone mass of young adult mice (7). This suggests that locally produced PTHrP is essential for normal decidualization, and through these actions influences embryonic bone growth. This indicates an additional role for PTHrP in maternal physiology.

**Figure 7:**
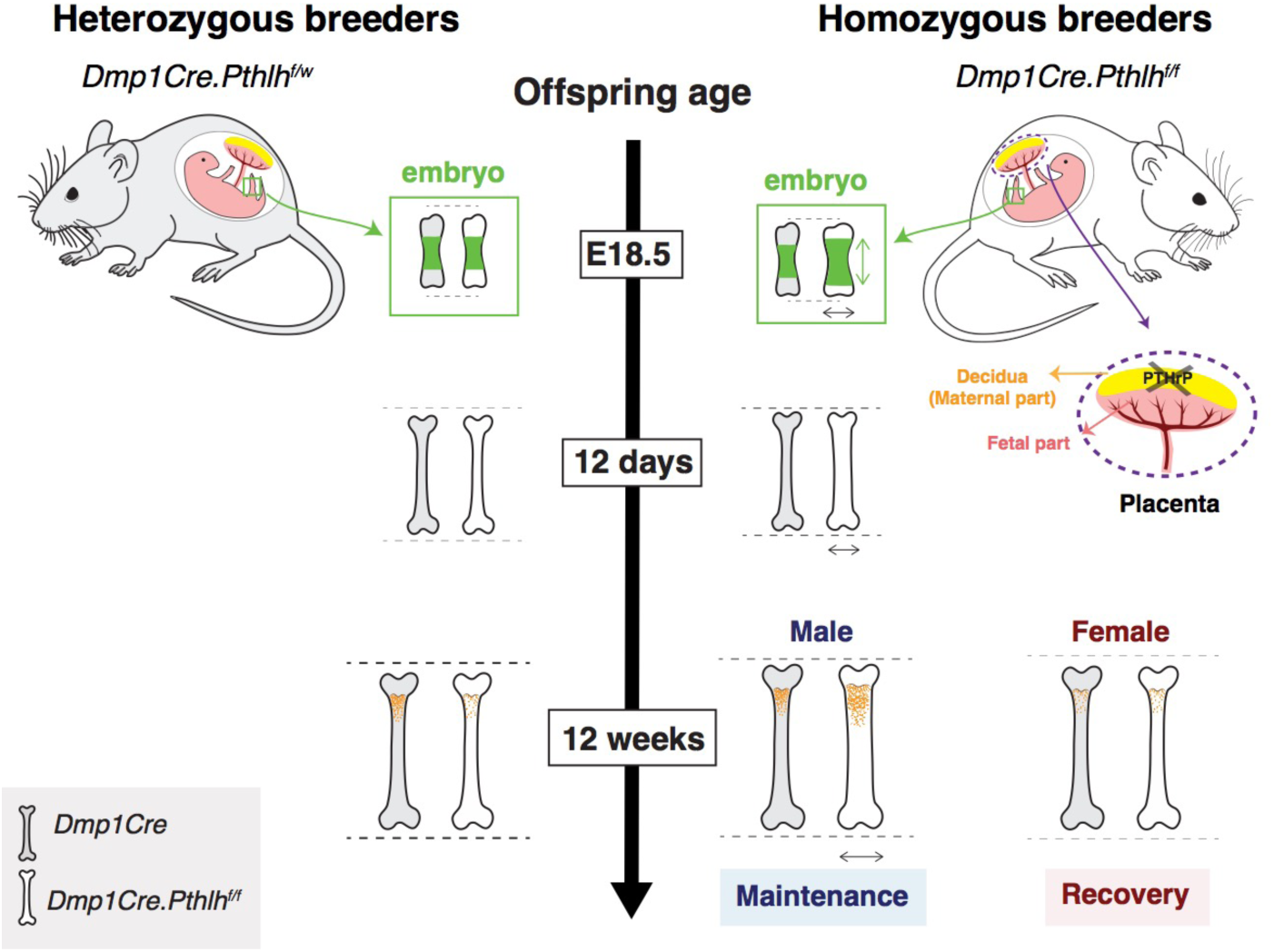
Decidual PTHrP determines bone width in progeny. Mice from breeders heterozygous for PTHrP (*Dmp1Cre.Pthlh*^*f/w*^ breeders) had normal bone size (length and width) compared to their sex- and age-matched controls, but lower adult trabecular bone mass. Decidual PTHrP may limit fetal skeletal development and radial growth, independent of longitudinal growth. Mothers of *Dmp1Cre.Pthlh*^*f/f(hom)*^ mice (which are *Dmp1Cre.Pthlh*^*f/f(het)*^) had lower levels of decidual PTHrP, leading to wider long bones in *Dmp1Cre.Pthlh*^*f/f(hom)*^ progeny. This phenotype was observed not only in embryos and neonatal mice, but also in adult male mice. Adult male mice also showed high trabecular bone mass compared to their sex-matched *Dmp1Cre* controls.

Our data contrasts the role of maternal-derived PTHrP with endogenous osteocytic PTHrP. *Dmp1Cre.Pthlh*^*f/f(hom)*^ mice exhibited wider long bones during late embryogenesis, at 12 days of age in both males and females, and during adulthood in males, leading to greater bone strength at 26 weeks of age. This phenotype was not observed in *Dmp1Cre.Pthlh*^*f/f(het)*^ mice at any time point studied. In addition to their increased bone width, adult male *Dmp1Cre.Pthlh*^*f/f(hom)*^ mice had high trabecular bone mass, in contrast to the osteopenia observed in *Dmp1Cre.Pthlh*^*f/f(het)*^ mice at 12 weeks of age (7). The requirement for decidual PTHrP therefore dominates the requirement for endogenous osteocytic PTHrP. The influence of decidual PTHrP in restricting bone width also contrasts with the role of endogenous PTHrP from the embryo-derived portion of the placenta, which promotes placental calcium transport and skeletal development, as indicated by reduced bone length in *Pthlh* null embryos (5).

The role of decidual PTHrP has not previously been investigated. PTHrP is produced within the decidua, myometrium, amnion, and chorion, where it dilates the uterine and placental vasculature and inhibits myometrial contractions (13-15). Here we showed PTHrP is expressed by decidua and spongiotrophoblast layer (or junctional zone), whereas labyrinth zone of placenta had undetectable levels of PTHrP. Decidua from mothers of *Dmp1Cre.Pthlh*^*f/f(hom)*^ mice had significantly lower PTHrP staining, but there was no change in placental PTHrP staining, supporting a maternal origin of the defect. Decidua from mothers of *Dmp1Cre.Pthlh*^*f/f(hom)*^ mice were also smaller in size and decidual cell morphology was modified. The smaller decidua and reduced size of decidual cells in mothers of *Dmp1Cre.Pthlh*^*f/f(hom)*^ mice was a surprising finding in this study. This may reflect reduced decidual cell differentiation, since decidual cells are enlarged during decidualization (16, 17). Alternatively, it may reflect decidual cell atrophy in mothers of *Dmp1Cre.Pthlh*^*f/f(hom)*^ mice. Since previous studies reported that PTHrP repressed decidualization of human uterine fibroblast cells (18), that intrauterine injection of PTH/PTHrP receptor antagonist from day 6 to 13 post coitum (after induction of decidua) increased decidualisation in rats (19), and *Pthlh* mRNA levels are downregulated during decidualization in primary endometrial stromal cells (20), we suggest the reduced size of decidual cells at day E17.5 may reflect early commencement of atrophy. How this drives the increased radial growth and mineralization in the *Dmp1Cre.Pthlh*^*f/f(hom)*^ embryo remains to be established.

It is very surprising that *Dmp1-Cre* targeted recombination had an influence on decidua. Although the *Dmp1-Cre* mouse is widely used as an osteocyte and late-osteoblast conditional knockout mouse, multiple off-target tissues have been reported. These include our previous report of recombination in the mammary gland (7). We and others have shown recombination in skeletal muscle and certain brain cells (7), and reporter genes have also shown *Dmp1-Cre* expression in preosteoblasts, a subset of bone marrow stromal cells, and gastrointestinal mesenchymal stromal cells (21). To date, there is no report that *Dmp1-Cre* targets decidua, which is a transient uterine tissue. We previously tested non-pregnant uterus, and found that *Dmp1-Cre* recombination did not occur (7). The expression of *Dmp1-Cre* in decidua has major implications for the design and reporting of experiments utilizing *Dmp1Cre* for gene deletion. However, this clearly depends on the function of the targeted gene. For example, although gp130, and its inhibitor protein SOCS3, are expressed in murine decidua (22, 23), homozygous-bred *Dmp1Cre.gp130*^*f/f*^ mice (8) and *Dmp1Cre.Socs3*^*f/f*^ mice (9) showed phenotypes similar to that of heterozygous-bred mice of the same genotype (9, 24).

Although decidua and placenta provide nutrition to promote embryonic and placental weight gain *in utero* (25), and these effects influence adult health, including bone mass (26), there was no change in total embryo or placental weight in *Dmp1Cre.Pthlh*^*f/f(hom)*^ embryos. This suggests maternal contributors to general embryonic nutrition, such as uteroplacental blood flow and nutrient transport across the placenta controlled by (for example) growth hormone, IGF-I, insulin, and glucocorticoids, are unlikely to contribute to the phenotype we observe. The influence of decidual PTHrP on growth appears to be specific to the skeleton. The alteration in the morphological features of decidua might be associated with changes in its function, resulting in changes in skeletal development. Whether decidual PTHrP, like embryonic placental PTHrP regulates placental calcium transport or fetal PTH levels (27, 28) or other local regulators of bone development, which are known to regulate fetal bone development (29), or it acts systematically to regulate bone radial growth remains unknown.

The increased mineralization length and greater bone width of *Dmp1Cre.Pthlh*^*f/f(hom)*^ embryos and adult male *Dmp1Cre.Pthlh*^*f/f(hom)*^ mice along the full length of the bone suggests that the maternal influence on bone widening determines radial expansion of the embryonic cartilage anlagen, the early cartilage model of the developing bone, rather than inducing periosteal apposition at the diaphysis. Maternal PTHrP may therefore limit radial expansion of chondrocytes during cartilage anlage development, and may suppress signalling pathways that promote expansion of chondrocytes at growth plate. This effect contrasts with actions of embryo-derived PTHrP from the placenta (5) and cartilage (30, 31) to stimulate longitudinal bone growth. Although longitudinal bone growth has been studied widely, very little is known about signaling pathways orchestrating radial growth. There are two non-mutually-exclusive theories describing how bone radial width is determined: the “mechanostat” theory suggests that bone size and shape are adapt to mechanical strain (32, 33), while the “sizostat” theory suggests a set of genes regulates bone width to reach a pre-programmed setting (34). Although different genomic markers have been correlated with bone size and bone shape (35), no specific genes or molecular pathways have yet been described as major determinants of cortical bone diameter. Our data suggests that decidual PTHrP is a determinant for the cortical width sizostat.

Although both male and female *Dmp1Cre.Pthlh*^*f/f(hom)*^ mice had wider long bones at 12 days of age, this phenotype was retained through to adulthood only in males, suggesting that the mechanisms controlling continued radial growth and bone width are sex-dependent. Although placental nutrition has sexually dimorphic effects on embryo growth (36), we did not observe sex differences in this study until after 12 days, again emphasizing that the wide bone phenotype is unlikely to relate to placental nutrition. Post-pubertal sex differences in cortical diameter are common to all mammals (36-45), with females having narrower bones than males, however the molecular mechanisms driving this sexual dimorphism remain largely unknown; this mouse model may therefore shed new light on the mechanisms that contribute to this sexual dimorphism. The retention of this phenotype in males, but not females, suggests that hormonal changes at puberty in females may slow their radial growth. While most studies investigating sexual dimorphism in murine bone width have focused on periosteal growth at the diaphysis, our results suggest that differences between males and females in cortical width might also arise from radial expansion of the growth plate. Estradiol is known to slow longitudinal growth: a previous study has shown that ovariectomy increased tibial length and increased chondrocyte proliferation (46), and 17beta-estradiol treatment of 26-day-old female and male rats led to shorter tibial length and an early reduction in growth plate longitudinal width (46). Testosterone also affects chondrocytes: local injection of testosterone into the tibial epiphyseal growth plate of castrated growing male rats significantly increased epiphyseal growth plate length (47). Furthermore, while the perinatal testosterone surge is required for adult bone length, bone width is determined by post-pubertal testosterone (48). The cellular and intracellular pathways by which estradiol and/or testosterone differentially affect growth plate radial growth in control and *Dmp1Cre.Pthlh*^*f/f(hom)*^ mice remains to be investigated.

In *Dmp1Cre.Pthlh*^*f/f(hom)*^ mice, greater cortical width was not associated with greater total bone length. To our knowledge, this is the first evidence of changes in bone diameter independent of cortical thickness, longitudinal growth, and total body weight gain. All previously reported mouse models with changes in bone diameter also showed widespread skeletal development defects such as reduced bone length and width or altered cortical thickness (48-51). For example, mice lacking the endogenous nuclear localization sequence and C-terminus of PTHrP displayed retarded growth with lower body weight and total skeletal size at the age of 2 weeks (50), while Insulin-like growth factor I null (*Igf1*^*-/-*^) mice displayed smaller body size, shortened femoral length and reduced cortical thickness compared to wild type littermates (51, 52). Having altered radial, but not longitudinal, growth makes the *Dmp1Cre.Pthlh*^*f/f(hom)*^ mouse an appealing model for studying specific mechanisms underlying radial bone growth. Discovering such mechanisms might open new therapeutic avenues for improving cortical bone strength and reducing fracture risk in growing children and in adults. If there is no change in material content, bones with wider cortices are more resistant to fracture (34, 53). Indeed, adult male *Dmp1Cre.Pthlh*^*f/f(hom)*^ bones could withstand greater force, and experienced lower ultimate stress, before failure in three point bending tests compared to age- and sex-matched controls.

Although we detected PTHrP recombination in the mammary glands (7), milk PTHrP levels were not significantly modified, and the wide-bone phenotype predated the commencement of suckling, indicating that a change in mammary supply of PTHrP is unlikely to cause the wide-bone phenotype we observed in *Dmp1Cre.Pthlh*^*f/f(hom)*^ mice. We had thought that this may have been a possibility since we previously noted *Dmp1Cre-* driven PTHrP recombination in the mammary gland (7), and suckling pups from mice lacking PTHrP in the milk supply (*BLG-Cre/PTHrP*^*lox/–*^) had higher ash calcium content, indicating greater bone mass, compared to controls at day 12 of lactation (10). The normal levels of PTHrP in milk in mothers of *Dmp1Cre.Pthlh*^*f/f(hom)*^ mice suggests that the mammary cells expressing *Dmp1Cre* are not the mammary epithelial cells that secrete PTHrP to the milk (54), and are different to those targeted in the *BLG-Cre/PTHrP*^*lox/–*^ model (10, 55, 56). Another possibility is that the level of PTHrP recombination was too low in mammary tissues to modify milk PTHrP production.

In conclusion, decidual PTHrP limits trabecular bone mass, bone geometry and strength, not only of neonatal mice, but also of adult male mice. *Dmp1Cre.Pthlh*^*f/f(hom)*^ embryos had accelerated skeletal development, with more mineralized and wider femora at E18.5. Although this effect was observed in both males and females in neonates, it was retained through to adulthood only in male mice. This indicates that maternal PTHrP limits bone growth, and this has a life-long influence on bone mass, shape and strength in male progeny.

## Materials and Methods

### Animal experiments

*Dmp1Cre.Pthlh*^*f/f(het)*^ mice have been described previously (7) and were bred from *Dmp1Cre* (Tg(*Dmp1*-cre)^1Jqfe^) mice (containing the *Dmp1* 10-kb promoter region) provided by Lynda Bonewald (University of Kansas, Kansas City, USA) (57), and *Pthlh*-flox (*Pthlh*^tm1Ack^) mice by Andrew Karaplis (McGill University, Montreal) (58) with LoxP sites spanning *Pthlh* exon III (7).

Two breeding strategies were used in this study (Figure 1A). Initially, mice hemizygous for *Dmp1Cre* were crossed with *Pthlh*^*f/f*^ mice to generate *Dmp1Cre.Pthlh*^*f/w*^ breeders. These were used to generate hemizygous-bred PTHrP deficient (*Dmp1Cre.Pthlh*^*f/f(het)*^) and *Dmp1Cre* mice, as in our previous study (7). *Dmp1Cre.Pthlh*^*f/f(hom)*^ mice were generated from breeding pairs that were both *Dmp1Cre.Pthlh*^*f/f(het)*^. Cousin-matched *Dmp1Cre* mice were bred to generate homozygous-bred *Dmp1Cre* cousin controls (denoted *Dmp1Cre*^*(hom)*^). Adult mice were collected at 14 weeks (male only), and at 26 weeks (both male and female) after an *in vivo* microCT scan at 16 weeks of age. Sample sizes used were based on our previous studies; no explicit power analysis was used.

12 day old pups were collected from both homozygous and heterozygous breeders. For the milk collection, mice were mated at 6 weeks of age, and after the mice became pregnant for the first time, males were removed. At day 12 of the (first) lactation, dams were anesthetized and injected with 1 IU of oxytocin (Sigma) (59). After 5 minutes, milk was collected and then kept at -80°C.

*Dmp1Cre.Pthlh*^*f/f(hom)*^ and *Dmp1Cre*^*(hom)*^ embryos and matching placenta/decidua were collected (4-7 litters/genotype) at embryonic day (E)18.5 of first pregnancy, and *Dmp1Cre.Pthlh*^*f/f(het)*^ embryos, placenta and decidua were collected at E17.5 of first pregnancy. The sex of embryos was determined by PCR, as described previously (60). All mice were housed at the St Vincent’s BioResources Centre, in a 12 h light and dark cycle and provided food and water *ad libitum*. St. Vincent’s Health Animal Ethics Committee approved all animal procedures. Terminal blood samples were collected by cardiac puncture exsanguination and sera kept at -80°C.

### Micro-computed tomography (micro-CT)

Micro-CT was carried out on samples from E17.5, E18.5, 12 days, 14 and 26 weeks of age. The observer was blinded to genotype and sex of all samples at the time of analysis. 26 week old mice were also anaesthetized and scanned by *in vivo* micro-CT at 16 weeks of age. After collection, embryos were fixed in 95% ethanol for 5 days. Femora of 14 and 26 week old mice, and tibiae of 12 day old mice were fixed overnight in 4% paraformaldehyde at 4°C, then stored in 70% ethanol until further analysis. Femoral and tibial morphology and microarchitecture were assessed using the Skyscan 1076 (E18.5, 12 days, 14 and 26 weeks of age) or 1276 (E17.5) micro-CT system (Bruker, Aartselaar, Belgium), as described previously (61) with the following modifications.

For micro-CT analysis at E17.5 and E18.5, embryos were scanned at 55 kV and 200 mA, and 48 kV and 208 mA, respectively. Projections were acquired over a pixel size of 5μm and 9 μm, respectively. Image slices were then reconstructed by NRecon (Bruker, version 1.7.1.0) with beam-hardening correction of 35%, ring artifact correction of 6, smoothing of 1, and defect pixel masking of 50%. The length of mineralized bone was measured in each femur. Femoral cortical structure was analyzed at three sites, based on the extent of mineralised femur: i) 20% of the mineralized length distal to the proximal end of the mineralized region (metaphysis; Top 20%); ii) Midshaft (Mid); iii) 20% of the mineralized length proximal to the distal end of the mineralized region (Bottom 20%). Automatic adaptive thresholding was used for each sample.

Tibiae from 12 day old mice were scanned at 37 kV and 228 mA. Regions of interest (ROI) commenced at a distance equal to 30% of the tibial length down the growth plate and an ROI of 10% of the tibial length was analyzed. The lower adaptive threshold limit used for cortical analysis was equivalent to 0.58 g/mm^3^ Calcium hydroxyapatite (CaHA).

Femora from 14, 16 and 26 week old mice were scanned at 45 kV and 220 mA. For trabecular and cortical analyses, ROI commenced at a distance equal to 7.5% or 30%, respectively, of the total femur length proximal to the distal end of the femur; for each, an ROI of 15% of the total femur length was analyzed. For 14 week old mice, the lower adaptive threshold limits for trabecular and cortical analysis were equivalent to 0.34 g/mm^3^ and 0.75 g/mm^3^ CaHA, respectively. For 16 week old mice, the lower adaptive threshold for trabecular and cortical analysis were equivalent to 0.30 g/mm^3^ and 0.64 g/mm^3^ CaHA, respectively. For 26 week old mice, the lower adaptive threshold for trabecular and cortical analysis were equivalent to 0.33 g/mm^3^ and 0.76 g/mm^3^ CaHA, respectively. For trabecular analysis in the 5^th^ lumbar vertebrae (L5), an ROI of half the height of the bone (vertically centered) with a diameter 2/3 the width of the vertebral body was analysed.

### Histomorphometry

Tibiae from 14 week old mice were embedded in methylmethacrylate and sectioned at 5 μm thickness for histomorphometric analysis, as previously described (62). The observer was blinded to genotype and sex of all samples during analysis. To determine bone formation rates, calcein was injected intraperitoneally (20 mg/kg) at 7 and 2 days before tissue collection. Sections were stained with Toluidine blue or Xylenol orange, as described (63). Static and dynamic histomorphometry of trabecular bone surfaces was carried out in the secondary spongiosa of the proximal tibia using the OsteoMeasure system (Osteometrics Inc., USA).

### Three-point bending test

Mechanical properties of femora were derived from three-point bending tests using a Bose Biodynamic 5500 Test Instrument (Bose, DE, USA), as described previously (64). The observer was blinded to genotype and sex of all samples during analysis. Once whole-bone properties were determined, tissue-level mechanical properties were calculated using micro-CT analysis of the mid-shaft (1).

### Biochemical assays

Cross-linked C-telopeptides of type I collagen (CTX-1) were measured in duplicate with the IDS RatLaps enzyme immunoassay (Abacus, Berkeley, CA, USA) in serum collected from mice fasted overnight. Serum levels of procollagen type 1 N propeptide (P1NP) were measured in duplicate using IDS Rat/Mouse PINP EIA kit (Abacus, Berkeley, CA, USA).

To measure milk PTHrP content, amino-terminal PTHrP RIA was carried out as previously described, with a sensitivity of 2 pM (65). Milk was diluted 1:500 in assay buffer prior to measurement of PTHrP. Milk PTHrP levels were also bioassayed as the cAMP generated in response to treatment of UMR106-01 cells, using PTH(1-34)-induced cAMP response as a standard curve (66). Replicate cell cultures in 24-well plates were incubated in cell culture medium with 1 mM isobutylmethylxanthine (IBMX) added. After treatment for 12 mins with 1:8 diluted milk samples, cAMP was measured by removing medium and adding acidified ethanol, drying, reconstituting in assay buffer and cAMP assay as described (67). cAMP was then corrected for total protein content of the milk, measured by Pierce BCA protein assay kit (Thermo Fisher Scientific). For this, milk was diluted 1:400 in PBS and absorbance was measured at OD562nm using the Polarstar Optima+ and a bovine serum albumin standard curve.

### Embryo skeletal staining

Alcian blue and Alizarin red S staining was carried out on E18.5 embryos, as described previously (68). Embryos were fixed in 95% ethanol for 5 days after skin removal. Remnant skin and viscera were dissected as much as possible, followed by defatting in acetone for 2 days. Thereafter, they were stained for 4 days at 40 °C in freshly prepared staining solution: 0.3% Alcian blue in 70% ethanol - 1 volume; 0.1% alizarin red S in 95% ethanol - 1volume; glacial acetic acid - 1 volume; 70% ethanol - 17 volumes. After washing in distilled water for 2 hours, they were cleared with 2% potassium hydroxide (KOH) for 2 days. Afterwards, they were put in 20% Glycerol in 1% KOH until skeletons were clearly visible, then successively placed into 50%, 80% and 100% glycerol solutions in 1% KOH for 2 days each. Femoral length was determined by measuring the distance between femoral head and distal end through a dissecting microscope, and an average of right and left femur lengths in each embryo was calculated.

### Immunohistochemistry

Immunohistochemistry was carried out as described previously (69, 70) on paraffin-embedded placenta/decidua (collected at E17.5) using goat rabbit anti-PTHrP (1:1000, R87, generated against PTHrP(1-14) (71). The observer was blinded to genotype and sex of all samples during analysis. Placental/decidual samples were fixed overnight in 4% paraformaldehyde at 4°C, stored in 70% ethanol, and embedded in paraffin wax until further analysis. Sections (5μm) were taken onto chrome alum-coated slides, dewaxed in Histoclear (National Diagnostics, Atlanta, GA), and rehydrated in graded ethanols. Endogenous peroxidase was blocked for 30 min in 2% H_2_O_2_ in methanol. After rinsing with 0.05 M phosphate buffered saline (PBS), samples were blocked with TNB (Renaissance TSA indirect (Tyramide Signal Amplification) PerkinElmer Life Sciences cat no-NEL700) for 60 min. PTHrP antibody (made against PTHrP (1-14) in house R87) 1:1000 or rabbit IgG (negative control) was applied for 2 hr at room temperature in a humid chamber. A secondary antibody (swine anti-rabbit, Dako) was applied for 30 min at 1:300, followed by streptavidin horseradish peroxidase (Dako) 1:300 in the same blocking solution for 30 min. PTHrP staining was visualized with diaminobenzidine kit (Dako) and conterstained with Mayer’s hematoxylin. Samples were rinsed in PBS between each step. PTHrP-DAB positive regions within the decidua and spongiotrophoblast zones were quantified with MetaMorph® image analysis software (Molecular Devices, San Jose, CA). The decidua and spongiotrophoblast zones were manually defined. Colour thresholding was applied and compared to negative controls. Integrated morphometry analysis was used to quantify DAB intensity, and frequency parameters.

### Statistical analysis

Statistically significant differences were determined by one-way or two-way ANOVA followed by Sidak’s post-hoc test or Fishers LSD posthoc test (uncorrected). Student’s t-test was used where only one comparison was being made. To analyse PTHrP positive areas in decidua/placentae, area under the curves were compared by Student’s t-test.

## Acknowledgments

The authors thank the staff of the St. Vincent’s Health Bioresources Centre for excellent animal care and assistance. This work was partially supported by National Health and Medical Research Council (Australia) Project Grants to N.A.S. and T.J.M. N.A. was supported by The University of Melbourne International Research Scholarship and a St. Vincent’s Institute top-up scholarship. T.I was supported by Travel Grants from Mochida Memorial Foundation for Medical and Pharmacological Research and The Foundation for Growth Science, Japan. N.A.S. is supported by a National Health and Medical Research Council (Australia) Senior Research Fellowship, and was supported in 2018 by the SVI Brenda Shanahan Fellowship. St. Vincent’s Institute is supported by the Victorian Government’s Operational Infrastructure Support Programme.

## Legends to Figure Supplements

**Figure 1 Supplement 1:**
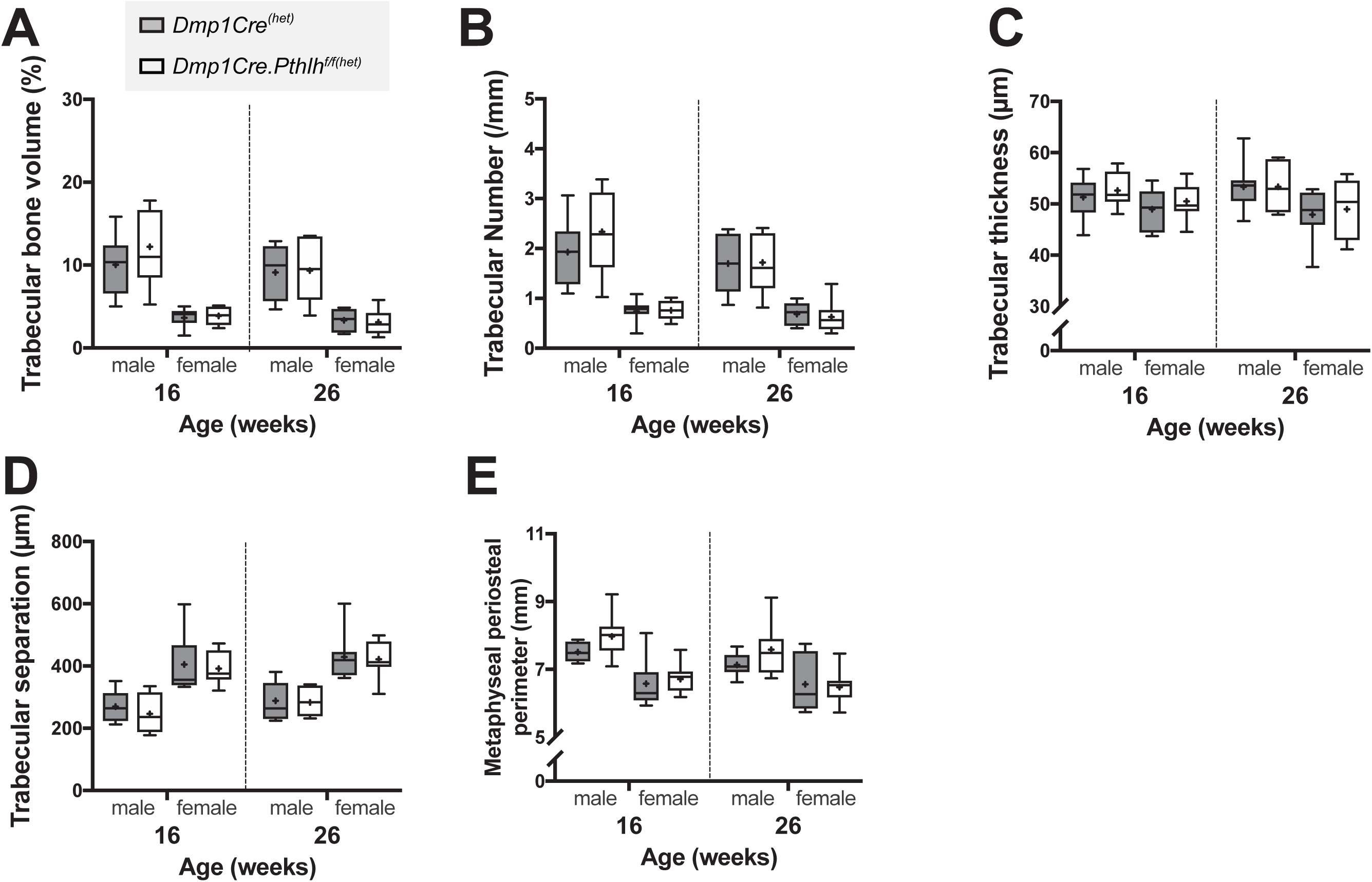
No difference in trabecular structure or metaphyseal diameter in femora from *Dmp1Cre.Pthlh*^*f/f(het)*^ mice and littermate *Dmp1Cre*^*(het)*^ controls. Trabecular structure of distal femoral primary spongiosa analysed by micro-CT in male and female mice at 16 and 26 weeks of age. Trabecular bone volume, trabecular number, trabecular thickness, trabecular separation, and metaphyseal periosteal perimeter are shown as mean (dot), interquartile range (box), median (line) and range; n=9-10/group. No significant differences associated with genotype were detected (two-way ANOVA). Y-axes are drawn to match those of Figure 1 to allow comparison with *Dmp1Cre.Pthlh*^*f/f(hom)*^ mice. Breeding strategy is shown in Figure 1A.

**Figure 2 Supplement 1:**
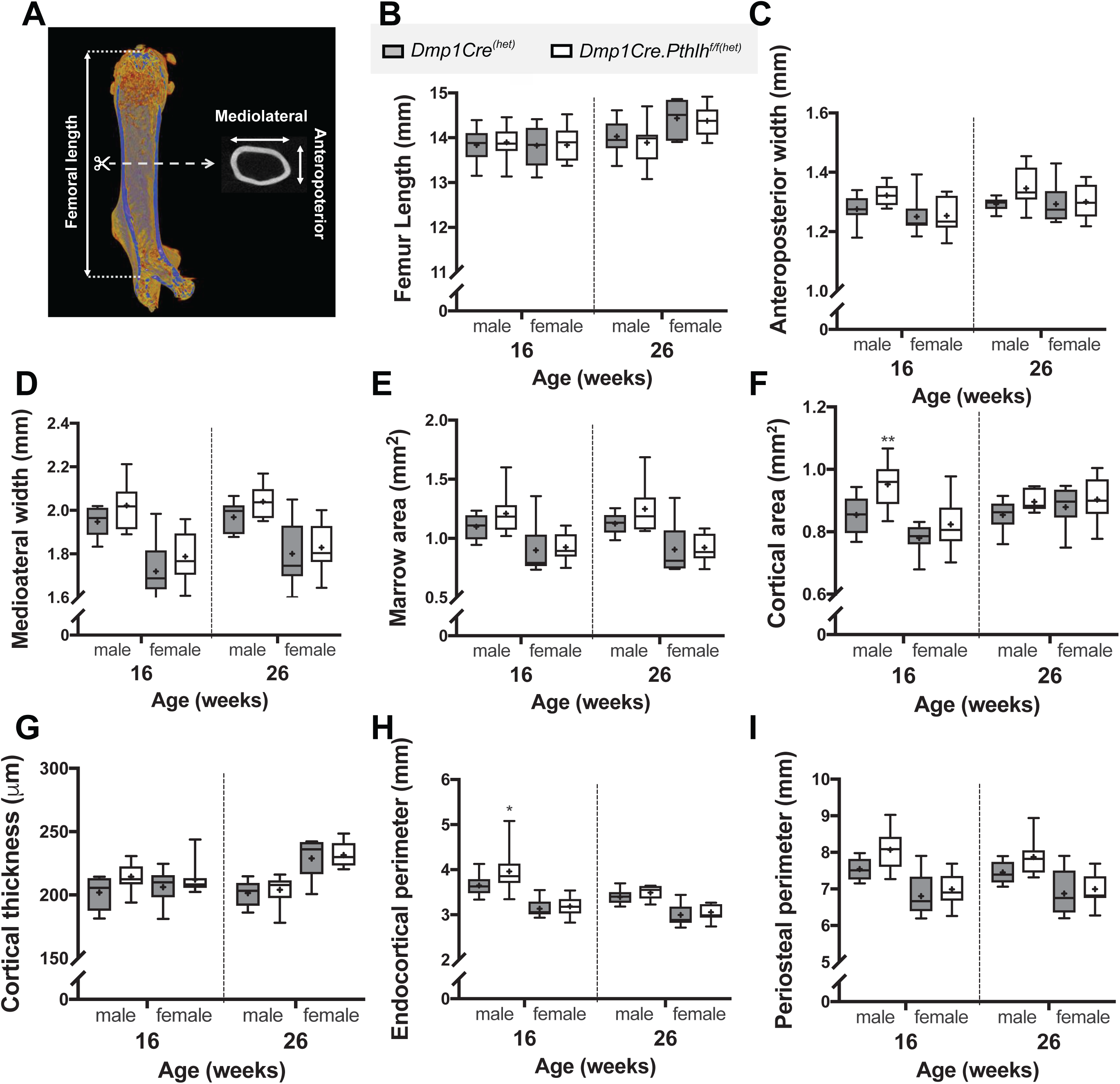
Cortical bone structure in femora from *Dmp1Cre.Pthlh*^*f/f(het)*^ mice and littermate *Dmp1Cre*^*(het)*^ controls. **A**: Schematic showing measurement regions (note, this is identical to Figure 2A, and is reproduced here for convenience). Length (**B**) and cortical dimensions of femora from male and female mice at 16 and 26 weeks of age. Anteroposterior (**C**) and mediolateral (**D**) width, measured by micro-CT at the midshaft. **E-I:** Femoral marrow area (**E**), cortical bone area (**F**), thickness (**G**), and both endocortical (**H**) and periosteal (**I**) perimeter were analysed in cortical ROI by micro-CT. Data are shown as mean (dot), interquartile range (box), median (line) and range; n=9-10/group. *p<0.05, and **p<0.01 compared to sex- and age-matched *Dmp1Cre*^*(het)*^ by two-way ANOVA. Y-axes are drawn to match those of Figure 2 to allow comparison with *Dmp1Cre.Pthlh*^*f/f(hom)*^ mice. Breeding strategy is shown in Figure 1A.

**Figure 3 Supplement 1:**
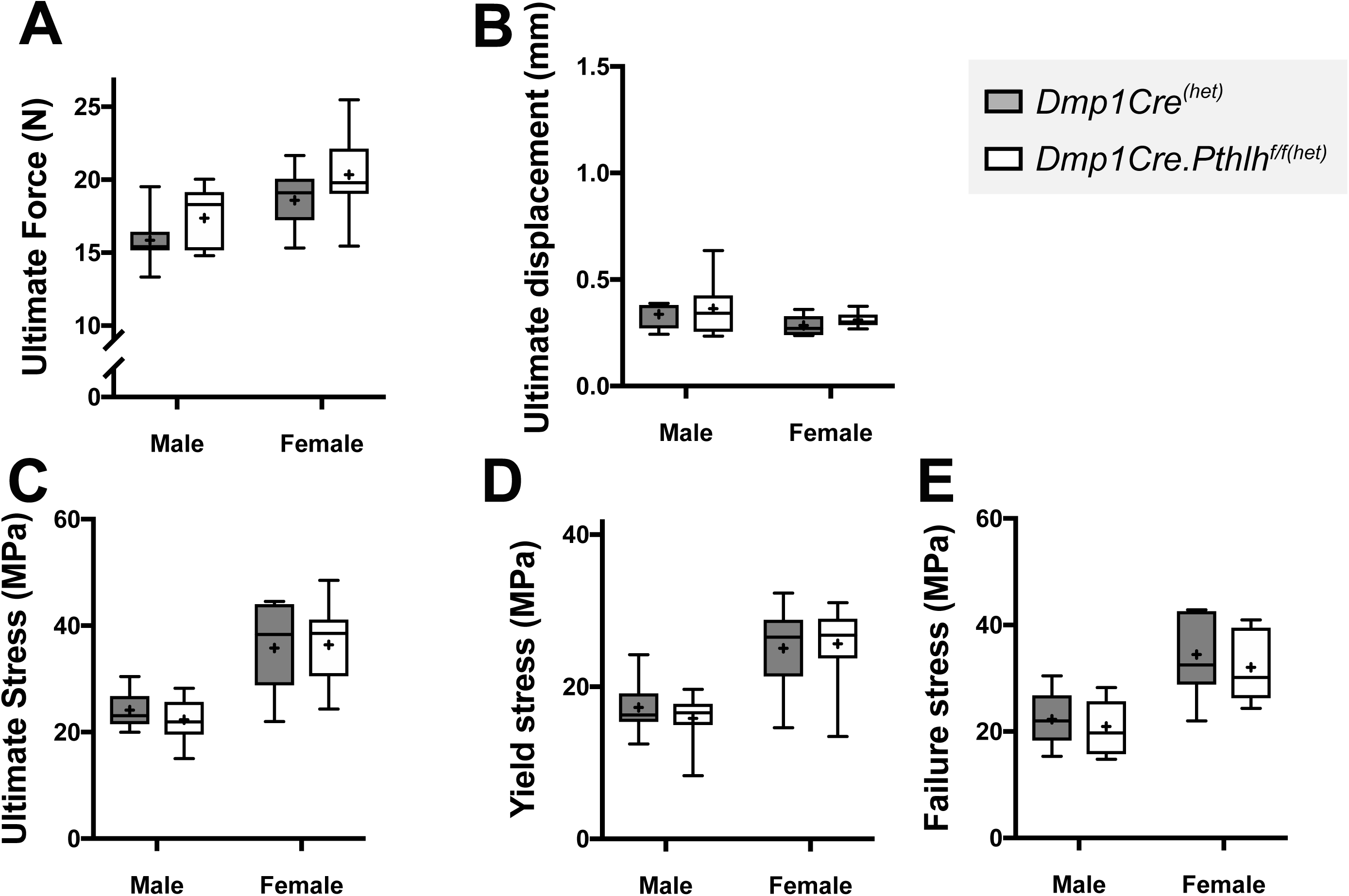
No change in bone strength measured by 3 point bending tests in heterozygous-bred *Dmp1Cre.Pthlh*^*f/f(het)*^ femora compared to littermate *Dmp1Cre*^*(het)*^ controls at 26 weeks of age. Shown are ultimate force (**A**), ultimate deformation (**B**), ultimate stress (**C**), yield stress (**D**), and failure stress (**E**). Data shown as mean (dot), interquartile range (box), median (line) and range, n=9-10/group. No significant differences relating to genotype were detected by two-way ANOVA.

**Figure 5 Supplement 1:**
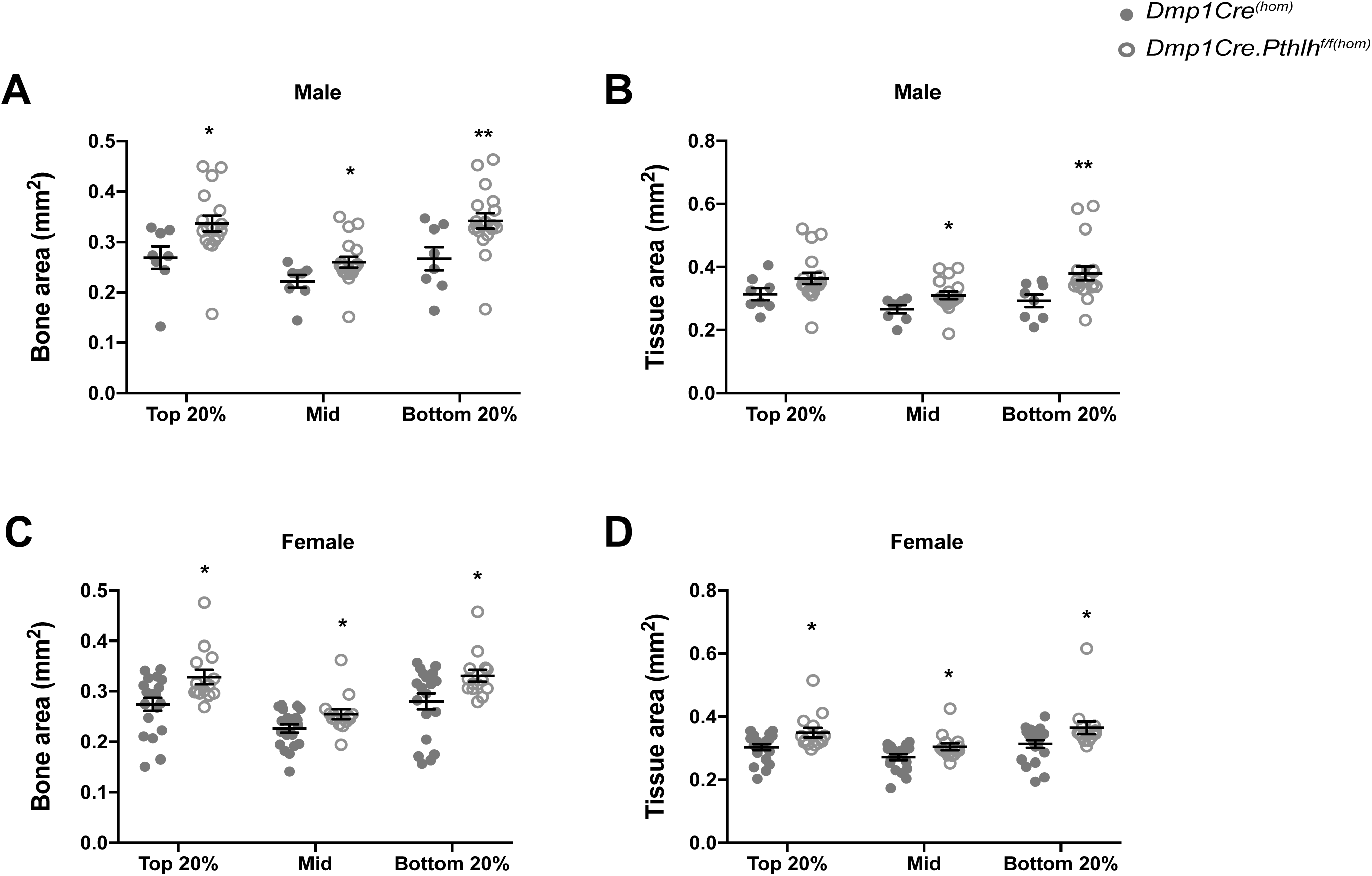
Additional micro-CT data of E18.5 *Dmp1Cre.Pthlh*^*f/f(hom)*^ embryos. Bone area and tissue area were analysed in cortical ROI by micro-CT in three different regions of femora: 20% of the mineralized length distal to the proximal end of the mineralized region (Top 20%), midshaft (Mid), and 20% of the mineralized length proximal to the distal end of the mineralized region (Bottom 20%); see Figure 5A. Data is shown as mean ± SEM with individual data points, *p<0.05 and **p<0.01 compared to controls by two-way ANOVA.

**Figure 5 Supplement 2:**
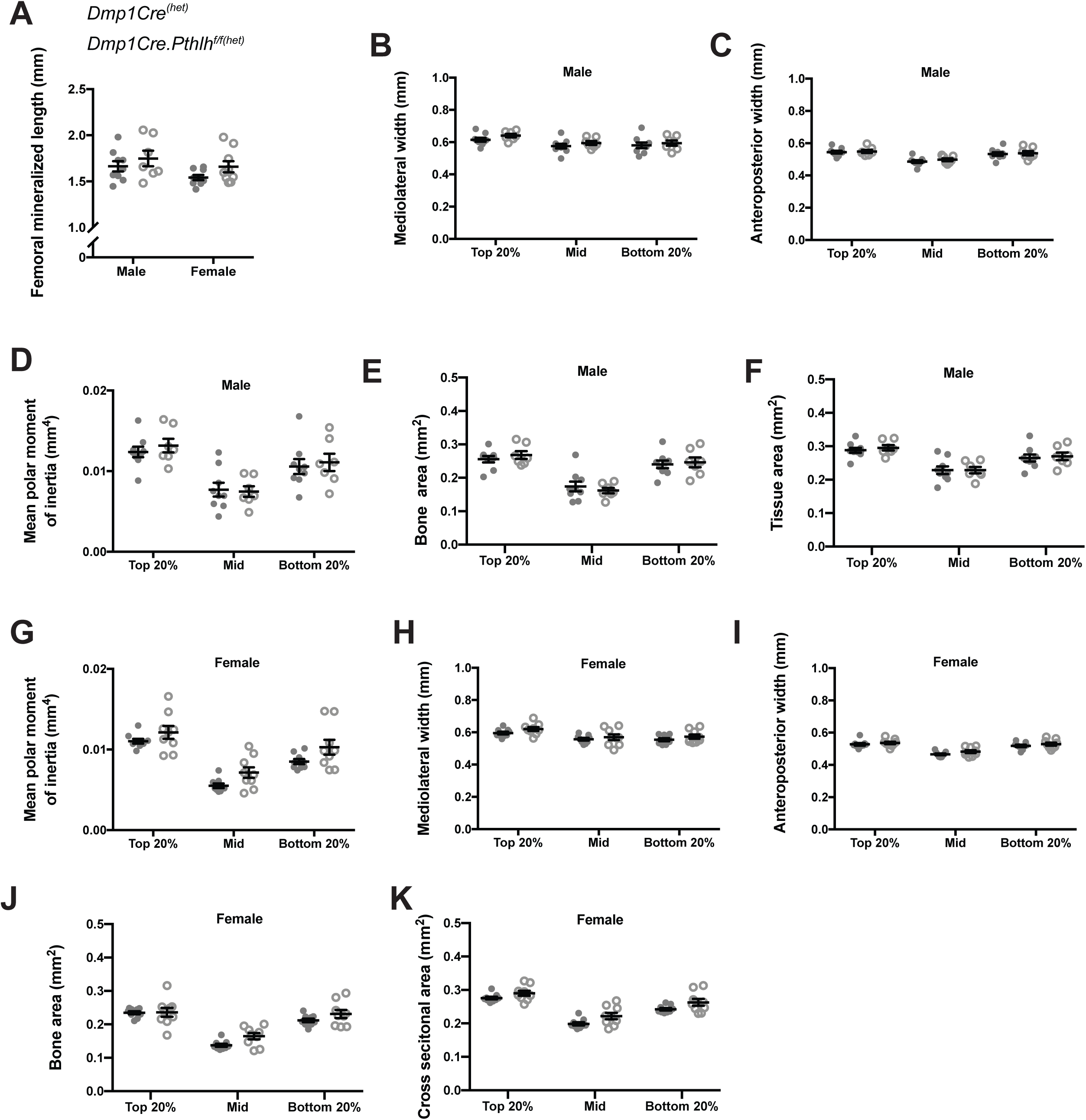
Bone structure of E17.5 *Dmp1Cre.Pthlh*^*f/f(het)*^ femora. **A**) Mineralized femoral length, mediolateral width (**B**,**H**), anteroposterior width (**C**,**I**), mean polar moment of inertia (**D**,**G**), bone area (**I**,**K**), and cross sectional area (**J**,**L**) of *Dmp1Cre.Pthlh*^*f/f(het)*^ and *Dmp1Cre*^*(het)*^ embryos measured in three different regions of femora: 20% of the mineralized length distal to the proximal end of the mineralized region (Top 20%), midshaft (Mid), and 20% of the mineralized length proximal to the distal end of the mineralized region (Bottom 20%). Data is shown as mean ± SEM with individual data points. No significant differences relating to genotype were detected by two-way ANOVA.

**Figure 6 Supplement 1:**
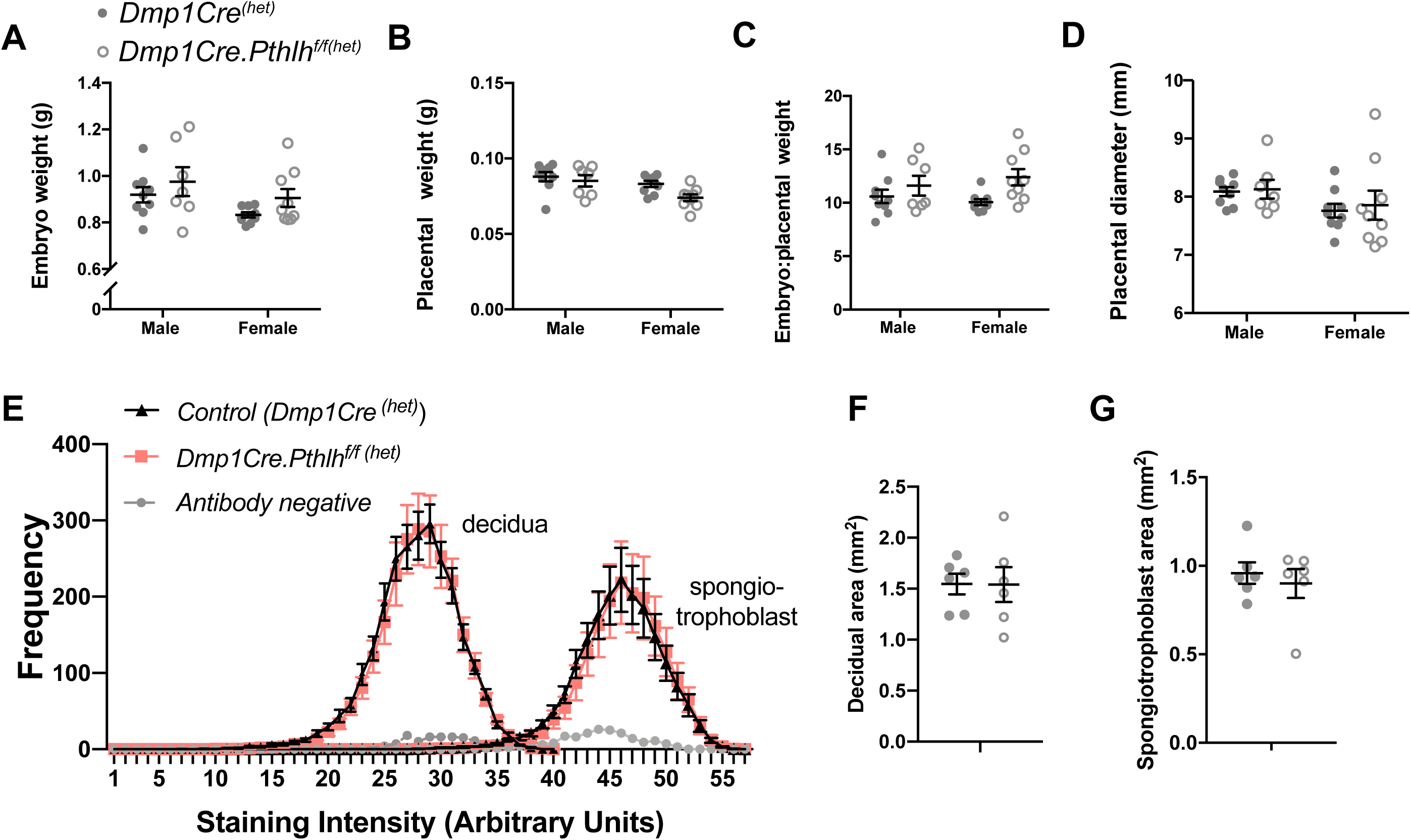
Embryo weight, placental weight, and decidual PTHrP of *Dmp1Cre.Pthlh*^*f/f(het)*^ mice. Embryo weight (**A**), placental weight (**B**) and embryo to placental weight ratio (**C**) of *Dmp1Cre.Pthlh*^*f/f(het)*^ and *Dmp1Cre*^*(het)*^ embryos at E17.5. (**D**) Frequency of PTHrP stained objects segregated by staining intensity in the spongiotrophoblast layer and decidua from placental/decidual samples from *Dmp1Cre.Pthlh*^*f/f(het)*^ and *Dmp1Cre*^*(het)*^ embryos. **(F**,**G)** Quantitation of total decidual area and spongiotrophoblast area; mean ± SEM with individual data points. No significant differences relating to genotype were detected by two-way ANOVA.

## References

1. Jepsen KJ, Silva MJ, Vashishth D, Guo XE, van der Meulen MC. Establishing biomechanical mechanisms in mouse models: practical guidelines for systematically evaluating phenotypic changes in the diaphyses of long bones. J Bone Miner Res. 2015;30(6):951–66.

2. Abad V, Meyers JL, Weise M, Gafni RI, Barnes KM, Nilsson O, et al. The role of the resting zone in growth plate chondrogenesis. Endocrinology. 2002;143(5):1851–7.

3. Pazzaglia UE, Beluffi G, Benetti A, Bondioni MP, Zarattini G. A review of the actual knowledge of the processes governing growth and development of long bones. Fetal and pediatric pathology. 2011;30(3):199–208.

4. Martin TJ. Parathyroid Hormone-Related Protein, Its Regulation of Cartilage and Bone Development, and Role in Treating Bone Diseases. Physiol Rev. 2016;96(3):831–71.

5. Karaplis AC, Luz A, Glowacki J, Bronson RT, Tybulewicz VL, Kronenberg HM, et al. Lethal skeletal dysplasia from targeted disruption of the parathyroid hormone-related peptide gene. Genes & Development. 1994;8(3):277–89.

6. Miao D, He B, Jiang Y, Kobayashi T, Soroceanu MA, Zhao J, et al. Osteoblast-derived PTHrP is a potent endogenous bone anabolic agent that modifies the therapeutic efficacy of administered PTH 1-34. J Clin Invest. 2005;115(9):2402–11.

7. Ansari N, Ho PW, Crimeen-Irwin B, Poulton IJ, Brunt AR, Forwood MR, et al. Autocrine and Paracrine Regulation of the Murine Skeleton by Osteocyte-Derived Parathyroid Hormone-Related Protein. J Bone Miner Res. 2018;33(1):137–53.

8. Standal T, Johnson RW, McGregor NE, Poulton IJ, Ho PW, Martin TJ, et al. gp130 in late osteoblasts and osteocytes is required for PTH-induced osteoblast differentiation. J Endocrinol. 2014;223(2):181–90.

9. Cho DC, Brennan HJ, Johnson RW, Poulton IJ, Gooi JH, Tonkin BA, et al. Bone corticalization requires local SOCS3 activity and is promoted by androgen action via interleukin-6. Nat Commun. 2017;8(1):806.

10. Mamillapalli R, VanHouten J, Dann P, Bikle D, Chang W, Brown E, et al. Mammary-specific ablation of the calcium-sensing receptor during lactation alters maternal calcium metabolism, milk calcium transport, and neonatal calcium accrual. Endocrinology. 2013;154(9):3031–42.

11. Karperien M, Lanser P, de Laat SW, Boonstra J, Defize LH. Parathyroid hormone related peptide mRNA expression during murine postimplantation development: evidence for involvement in multiple differentiation processes. The International journal of developmental biology. 1996;40(3):599–608.

12. Kovacs CS, Lanske B, Hunzelman JL, Guo J, Karaplis AC, Kronenberg HM. Parathyroid hormone-related peptide (PTHrP) regulates fetal-placental calcium transport through a receptor distinct from the PTH/PTHrP receptor. Proc Natl Acad Sci U S A. 1996;93(26):15233–8.

13. Ferguson JE, 2nd, Gorman JV, Bruns DE, Weir EC, Burtis WJ, Martin TJ, et al. Abundant expression of parathyroid hormone-related protein in human amnion and its association with labor. Proceedings of the National Academy of Sciences of the United States of America. 1992;89(17):8384–8.

14. Williams ED, Leaver DD, Danks JA, Moseley JM, Martin TJ. Effect of parathyroid hormone-related protein (PTHrP) on the contractility of the myometrium and localization of PTHrP in the uterus of pregnant rats. J Reprod Fertil. 1994;102(1):209–14.

15. Meziani F, Van Overloop B, Schneider F, Gairard A. Parathyroid hormone-related protein-induced relaxation of rat uterine arteries: influence of the endothelium during gestation. J Soc Gynecol Investig. 2005;12(1):14–9.

16. Ramathal CY, Bagchi IC, Taylor RN, Bagchi MK. Endometrial decidualization: of mice and men. Semin Reprod Med. 2010;28(1):17–26.

17. Okada H, Tsuzuki T, Murata H. Decidualization of the human endometrium. Reprod Med Biol. 2018;17(3):220–7.

18. Sherafat-Kazemzadeh R, Schroeder JK, Kessler CA, Handwerger S. Parathyroid hormone-like hormone (PTHLH) represses decidualization of human uterine fibroblast cells by an autocrine/paracrine mechanism. The Journal of clinical endocrinology and metabolism. 2011;96(2):509–14.

19. Williams ED, Major BJ, Martin TJ, Moseley JM, Leaver DD. Effect of antagonism of the parathyroid hormone (PTH)/PTH-related protein receptor on decidualization in rat uterus. J Reprod Fertil. 1998;112(1):59–67.

20. Katoh N, Kuroda K, Tomikawa J, Ogata-Kawata H, Ozaki R, Ochiai A, et al. Reciprocal changes of H3K27ac and H3K27me3 at the promoter regions of the critical genes for endometrial decidualization. Epigenomics. 2018;10(9):1243–57.

21. Lim J, Burclaff J, He G, Mills JC, Long F. Unintended targeting of. Bone Res. 2017;5:16049.

22. San Martin S, Fitzgerald JS, Weber M, Párraga M, Sáez T, Zorn TM, et al. Stat3 and Socs3 expression patterns during murine placenta development. Eur J Histochem. 2013;57(2):e19–e.

23. Ni H, Ding N-Z, Harper MJK, Yang Z-M. Expression of leukemia inhibitory factor receptor and gp130 in mouse uterus during early pregnancy. Molecular reproduction and development. 2002;63(2):143–50.

24. Johnson RW, Brennan HJ, Vrahnas C, Poulton IJ, McGregor NE, Standal T, et al. The primary function of gp130 signaling in osteoblasts is to maintain bone formation and strength, rather than promote osteoclast formation. J Bone Miner Res. 2014;29(6):1492–505.

25. Murphy VE, Smith R, Giles WB, Clifton VL. Endocrine regulation of human fetal growth: the role of the mother, placenta, and fetus. Endocrine reviews. 2006;27(2):141–69.

26. Holroyd CR, Osmond C, Barker DJ, Ring SM, Lawlor DA, Tobias JH, et al. Placental Size Is Associated Differentially With Postnatal Bone Size and Density. J Bone Miner Res. 2016;31(10):1855–64.

27. Smith FG, Jr., Alexander DP, Buckle RM, Britton HG, Nixon DA. Parathyroid hormone in foetal and adult sheep: the effect of hypocalcaemia. The Journal of endocrinology. 1972;53(3):339–48.

28. Brown EM, Gamba G, Riccardi D, Lombardi M, Butters R, Kifor O, et al. Cloning and characterization of an extracellular Ca(2+)-sensing receptor from bovine parathyroid. Nature. 1993;366(6455):575–80.

29. Miao D, He B, Karaplis AC, Goltzman D. Parathyroid hormone is essential for normal fetal bone formation. J Clin Invest. 2002;109(9):1173–82.

30. Lanske B, Karaplis AC, Lee K, Luz A, Vortkamp A, Pirro A, et al. PTH/PTHrP receptor in early development and Indian hedgehog-regulated bone growth. Science. 1996;273(5275):663–6.

31. Kronenberg HM. PTHrP and skeletal development. Annals of the New York Academy of Sciences. 2006;1068:1–13.

32. Frost HM. Mechanical determinants of bone modeling. Metab Bone Dis Relat Res. 1982;4(4):217–29.

33. Frost HM. The mechanostat: a proposed pathogenic mechanism of osteoporoses and the bone mass effects of mechanical and nonmechanical agents. Bone Miner. 1987;2(2):73–85.

34. Rauch F. Bone growth in length and width: the Yin and Yang of bone stability. J Musculoskelet Neuronal Interact. 2005;5(3):194–201.

35. Volkman SK, Galecki AT, Burke DT, Paczas MR, Moalli MR, Miller RA, et al. Quantitative trait loci for femoral size and shape in a genetically heterogeneous mouse population. Journal of bone and mineral research : the official journal of the American Society for Bone and Mineral Research. 2003;18(8):1497–505.

36. Aiken CE, Ozanne SE. Sex differences in developmental programming models. Reproduction. 2013;145(1):R1–13.

37. Oyster N. Sex differences in cancellous and cortical bone strength, bone mineral content and bone density. Age Ageing. 1992;21(5):353–6.

38. Nelson DA, Megyesi MS. Sex and ethnic differences in bone architecture. Current osteoporosis reports. 2004;2(2):65–9.

39. Gabel L, Macdonald HM, McKay HA. Sex Differences and Growth-Related Adaptations in Bone Microarchitecture, Geometry, Density, and Strength From Childhood to Early Adulthood: A Mixed Longitudinal HR-pQCT Study. Journal of bone and mineral research : the official journal of the American Society for Bone and Mineral Research. 2017;32(2):250–63.

40. Evans RK, Negus C, Antczak AJ, Yanovich R, Israeli E, Moran DS. Sex differences in parameters of bone strength in new recruits: beyond bone density. Med Sci Sports Exerc. 2008;40(11 Suppl):S645–53.

41. Eckstein F, Matsuura M, Kuhn V, Priemel M, Muller R, Link TM, et al. Sex differences of human trabecular bone microstructure in aging are site-dependent. J Bone Miner Res. 2007;22(6):817–24.

42. Beck TJ, Ruff CB, Scott WW, Jr., Plato CC, Tobin JD, Quan CA. Sex differences in geometry of the femoral neck with aging: a structural analysis of bone mineral data. Calcified tissue international. 1992;50(1):24–9.

43. Amin S, Khosla S. Sex- and age-related differences in bone microarchitecture in men relative to women assessed by high-resolution peripheral quantitative computed tomography. J Osteoporos. 2012;2012:129760.

44. Sigurdsson G, Aspelund T, Chang M, Jonsdottir B, Sigurdsson S, Eiriksdottir G, et al. Increasing sex difference in bone strength in old age: The Age, Gene/Environment Susceptibility-Reykjavik study (AGES-REYKJAVIK). Bone. 2006;39(3):644–51.

45. Ackerman A, Thornton JC, Wang J, Pierson RN, Jr., Horlick M. Sex difference in the effect of puberty on the relationship between fat mass and bone mass in 926 healthy subjects, 6 to 18 years old. Obesity (Silver Spring). 2006;14(5):819–25.

46. van der Eerden BC, Emons J, Ahmed S, van Essen HW, Lowik CW, Wit JM, et al. Evidence for genomic and nongenomic actions of estrogen in growth plate regulation in female and male rats at the onset of sexual maturation. The Journal of endocrinology. 2002;175(2):277–88.

47. Ren SG, Malozowski S, Sanchez P, Sweet DE, Loriaux DL, Cassorla F. Direct administration of testosterone increases rat tibial epiphyseal growth plate width. Acta Endocrinol (Copenh). 1989;121(3):401–5.

48. Sims NA, Brennan K, Spaliviero J, Handelsman DJ, Seibel MJ. Perinatal testosterone surge is required for normal adult bone size but not for normal bone remodeling. Am J Physiol Endocrinol Metab. 2006;290(3):E456–62.

49. Ye L, Mishina Y, Chen D, Huang H, Dallas SL, Dallas MR, et al. Dmp1-deficient mice display severe defects in cartilage formation responsible for a chondrodysplasia-like phenotype. The Journal of biological chemistry. 2005;280(7):6197–203.

50. Miao D, Su H, He B, Gao J, Xia Q, Zhu M, et al. Severe growth retardation and early lethality in mice lacking the nuclear localization sequence and C-terminus of PTH-related protein. Proc Natl Acad Sci U S A. 2008;105(51):20309–14.

51. Bikle D, Majumdar S, Laib A, Powell-Braxton L, Rosen C, Beamer W, et al. The skeletal structure of insulin-like growth factor I-deficient mice. Journal of bone and mineral research : the official journal of the American Society for Bone and Mineral Research. 2001;16(12):2320–9.

52. Liu JP, Baker J, Perkins AS, Robertson EJ, Efstratiadis A. Mice carrying null mutations of the genes encoding insulin-like growth factor I (Igf-1) and type 1 IGF receptor (Igf1r). Cell. 1993;75(1):59–72.

53. Vashishth D. Small animal bone biomechanics. Bone. 2008;43(5):794–7.

54. Ferrari SL, Rizzoli R, Bonjour JP. Parathyroid hormone-related protein production by primary cultures of mammary epithelial cells. Journal of cellular physiology. 1992;150(2):304–11.

55. Hyttinen JM, Korhonen VP, Hiltunen MO, Myohanen S, Janne J. High-level expression of bovine beta-lactoglobulin gene in transgenic mice. J Biotechnol. 1998;61(3):191–8.

56. VanHouten JN, Dann P, Stewart AF, Watson CJ, Pollak M, Karaplis AC, et al. Mammary-specific deletion of parathyroid hormone–related protein preserves bone mass during lactation. Journal of Clinical Investigation. 2003;112(9):1429–36.

57. Lu Y, Xie Y, Zhang S, Dusevich V, Bonewald LF, Feng JQ. DMP1-targeted Cre expression in odontoblasts and osteocytes. J Dent Res. 2007;86(4):320–5.

58. He B, Deckelbaum RA, Miao D, Lipman ML, Pollak M, Goltzman D, et al. Tissue-specific targeting of the pthrp gene: the generation of mice with floxed alleles. Endocrinology. 2001;142(5):2070–7.

59. Gillies BR, Ryan BA, Tonkin BA, Poulton IJ, Ma Y, Kirby BJ, et al. Absence of Calcitriol Causes Increased Lactational Bone Loss and Lower Milk Calcium but Does Not Impair Post-lactation Bone Recovery in Cyp27b1 Null Mice. J Bone Miner Res. 2018;33(1):16–26.

60. McFarlane L, Truong V, Palmer JS, Wilhelm D. Novel PCR assay for determining the genetic sex of mice. Sex Dev. 2013;7(4):207–11.

61. Takyar FM, Tonna S, Ho PW, Crimeen-Irwin B, Baker EK, Martin TJ, et al. EphrinB2/EphB4 inhibition in the osteoblast lineage modifies the anabolic response to parathyroid hormone. J Bone Miner Res. 2013;28(4):912–25.

62. Sims NA, Clement-Lacroix P, Da Ponte F, Bouali Y, Binart N, Moriggl R, et al. Bone homeostasis in growth hormone receptor-null mice is restored by IGF-I but independent of Stat5. J Clin Invest. 2000;106(9):1095–103.

63. Sims NA, Jenkins BJ, Nakamura A, Quinn JM, Li R, Gillespie MT, et al. Interleukin-11 receptor signaling is required for normal bone remodeling. J Bone Miner Res. 2005;20(7):1093–102.

64. Williamson L, Hayes A, Hanson ED, Pivonka P, Sims NA, Gooi JH. High dose dietary vitamin D3 increases bone mass and strength in mice. Bone Rep. 2017;6:44–50.

65. Grill V, Ho P, Body JJ, Johanson N, Lee SC, Kukreja SC, et al. Parathyroid hormone-related protein: elevated levels in both humoral hypercalcemia of malignancy and hypercalcemia complicating metastatic breast cancer. J Clin Endocrinol Metab. 1991;73(6):1309–15.

66. Ho PW, Goradia A, Russell MR, Chalk AM, Milley KM, Baker EK, et al. Knockdown of PTHR1 in osteosarcoma cells decreases invasion and growth and increases tumor differentiation in vivo. Oncogene. 2015;34(22):2922–33.

67. Hammonds RG, Jr., McKay P, Winslow GA, Diefenbach-Jagger H, Grill V, Glatz J, et al. Purification and characterization of recombinant human parathyroid hormone-related protein. The Journal of biological chemistry. 1989;264(25):14806–11.

68. McLeod MJ. Differential staining of cartilage and bone in whole mouse fetuses by alcian blue and alizarin red S. Teratology. 1980;22(3):299–301.

69. McGregor NE, Poulton IJ, Walker EC, Pompolo S, Quinn JM, Martin TJ, et al. Ciliary neurotrophic factor inhibits bone formation and plays a sex-specific role in bone growth and remodeling. Calcif Tissue Int. 2010;86(3):261–70.

70. Walker EC, McGregor NE, Poulton IJ, Solano M, Pompolo S, Fernandes TJ, et al. Oncostatin M promotes bone formation independently of resorption when signaling through leukemia inhibitory factor receptor in mice. J Clin Invest. 2010;120(2):582–92.

71. Lam MH, Thomas RJ, Loveland KL, Schilders S, Gu M, Martin TJ, et al. Nuclear transport of parathyroid hormone (PTH)-related protein is dependent on microtubules. Molecular endocrinology. 2002;16(2):390–401.

